# Glutathione binding to the plant *At*Atm3 transporter and implications for the conformational coupling of ABC transporters

**DOI:** 10.1101/2021.12.13.472443

**Authors:** Chengcheng Fan, Douglas C. Rees

## Abstract

The ATP Binding Cassette (ABC) transporter of mitochondria (Atm) from *Arabidopsis thaliana* (*At*Atm3) has been implicated in the maturation of cytosolic iron-sulfur proteins and heavy metal detoxification, plausibly by exporting glutathione derivatives. Using single-particle cryo-electron microscopy, we have determined structures of *At*Atm3 in multiple conformational states. These structures not only provide a structural framework for defining the alternating access transport cycle, but also highlight an unappreciated feature of the glutathione binding site, namely the paucity of cysteine residues that could potentially form inhibitory mixed disulfides with glutathione. Despite extensive efforts, we were unable to prepare the ternary complex of *At*Atm3 with bound GSSG and MgATP. A survey of structurally characterized type IV ABC transporters that includes *At*Atm3 establishes that while nucleotides are found associated with all conformational states, they are effectively required to stabilize occluded and outward-facing conformations. In contrast, transport substrates have only been observed associated with inward-facing conformations. The absence of structures containing both nucleotide and transport substrate suggests that this ternary complex exists only transiently during the transport cycle.

## Background

The ATP Binding Cassette (ABC) transporter of mitochondria (Atm) family plays a vital (Leighton and Schatz, 1995), but enigmatic, role broadly related to transition metal homeostasis in eukaryotes (Lill et al., 2014). The best characterized member is *Saccharomyces cerevisiae* Atm1 (*Sc*Atm1) present in the inner membrane of mitochondria (Leighton and Schatz, 1995) and required for formation of cytosolic iron-sulfur cluster containing proteins (Kispal et al., 1999). Defects in *Sc*Atm1 lead to an overaccumulation of iron in the mitochondria (Kispal et al., 1997). Atm1 is proposed to transport a sulfur containing intermediate (Kispal et al., 1999) that may also include iron (Pandey et al., 2019). It is also likely to transport a similar sulfur containing species from the mitochondria that is required for the cytoplasmic thiolation of tRNA (Pandey et al., 2018). While the precise substrate that is transported remains unknown, derivatives of glutathione have been implicated based on their ability to stimulate the ATPase activity of Atm1 (Kuhnke et al., 2006).

Structures for Atm family members are currently available for *Sc*Atm1 (Srinivasan et al., 2014), the bacterial homolog *Na*Atm1 from *Novosphingobium aromaticivorans* (Lee et al., 2014) and human ABCB6 (Wang et al., 2020); the pairwise sequence identities between these homologous transporters range from 40% to 46%. These proteins occur as homodimers of half-transporters, where each half-transporter contains a transmembrane domain (TMD) followed by the canonical nucleotide binding domain (NBD) that defines the ABC transporter family. Each TMD consists of six transmembrane helices (TMs) that exhibit the exporter type I fold first observed for Sav1866 (Dawson and Locher, 2006); a recent re-classification now identifies this group as type IV ABC transporters (Thomas et al., 2020). The translocation of substrates across the membrane proceeds through an alternating access mechanism involving the ATP dependent interconversion between inward- and outward-facing conformational states. Among the Atm1 family, these conformations have been most extensively characterized for *Na*Atm1 and include the occluded and closed states that provide a structural framework for the unidirectional transport cycle (Fan et al., 2020). Structures of *Sc*Atm1 with reduced glutathione (GSH) (Srinivasan et al., 2014), and of *Na*Atm1 bound to reduced (GSH), oxidized (GSSG) and metallated (GS-Hg-SG) (Lee et al., 2014), have defined the general substrate binding site in the TMD for the transport substrates.

Plants have been found to have large numbers of transporters (Hwang et al., 2016), including *Arabidopsis* with three Atm orthologues, *At*Atm1, *At*Atm2, and *At*Atm3 (Chen et al., 2007). Of these, *At*Atm3 (also known as ABCB25) rescues the *Sc*Atm1 phenotype (Chen et al., 2007), and has been shown to be associated with maturation of cytosolic iron-sulfur proteins (Kushnir et al., 2001), confer resistance to heavy metals such as cadmium and lead (Kim et al., 2006), and participate in the formation of molybdenum-cofactor containing enzymes (Bernard et al., 2009; Teschner et al., 2010). Unlike yeast, defects in *At*Atm3 are not associated with iron accumulation in mitochondria (Bernard et al., 2009). While the physiological substrate is unknown, *At*Atm3 has been shown to transport oxidized glutathione and glutathione polysulfide (GSSSG), with the persulfidated species perhaps relevant to cytosolic iron-sulfur cluster assembly (Schaedler et al., 2014). The ability of *At*Atm3 to export oxidized glutathione has been implicated in helping stabilize against excessive glutathione oxidation in the mitochondria and thereby serving to maintain a suitable reduction potential (Marty et al., 2019).

To help address the functional role(s) of Atm transporters, we have determined structures of *At*Atm3 in multiple conformational states by single-particle cryo-electron microscopy (cryoEM). These structures not only provide a structural framework for defining the alternating access transport cycle, but also highlight an unappreciated feature of the glutathione binding site, namely the paucity of cysteine residues that could potentially form inhibitory mixed disulfides during the transport cycle. A survey of structurally characterized members of the type IV family of ABC transporters, including the Atm1 family, establishes that nucleotides are effectively required for the stabilization of the closed, occluded, and outward-facing conformations. In contrast to the nucleotide states, transport substrates and related inhibitors have only been observed associated with inward-facing conformational states. The absence of structures containing both nucleotide and transport substrate suggests that this ternary complex exists only transiently during the transport cycle.

## Results

*At*Atm3 contains an N-terminal mitochondrial targeting sequence that directs the translated protein to the mitochondria, where it is cleaved following delivery to the inner membrane. Since this targeting sequence consists of ∼80 residues and is anticipated to be poorly ordered, we generated three different N-terminal truncation mutants of *At*Atm3 through deletion of 60, 70 or 80 residues to identify the best-behaved construct. Together with the wild type construct, these three variants were heterologously overexpressed in *E. coli*. The construct with the 80 amino acids deletion showed the highest expression level and proportionally less aggregation by size exclusion chromatography (Figure S1) and hence was used for further functional and structural studies.

### ATPase activities

Using the 80-residue truncation construct, *At*Atm3 was purified in the detergent dodecyl-*β*-D-maltoside (DDM) and reconstituted into nanodiscs formed from the membrane scaffolding protein (MSP) 1D1 and the lipid 1-palmitoyl-2-oleoyl-glycero-3-phosphocholine (POPC). The ATPase activity of this construct was measured as a function of MgATP concentration in the absence and presence of either 2.5 mM GSSG or 10 mM GSH, which approximate their physiological concentrations in *E. coli* (Bennett et al., 2009). The rate of ATP hydrolysis was determined by measuring phosphate release using a molybdate based colorimetric ATPase activity assay (Chifflet, 1988). The basal ATPase activity, measured in the absence of glutathione derivatives, was significantly higher in detergent than in nanodiscs (104 versus 7.7 nmol min^-1^ mg^-1^, respectively; Figures 1ab), while the apparent K_m_s for MgATP were within a factor of two (∼0.16 mM and 0.08 mM, respectively). The ATPase activity of *At*Atm3 is stimulated by both 2.5 mM GSSG and 10 mM GSH, but the extent of stimulation depends strongly on the solubilization conditions. In nanodiscs, the ATPase rates increase to 32 and 39 nmol min^-1^ mg^-1^ with 2.5 mM GSSG and 10 mM GSH, respectively, for an overall increase of 4-5x above the basal rate. The ATPase rates for *At*Atm3 in DDM also increase with GSSG and GSH, to 117 and 154 nmol min^-1^ mg^-1^, respectively. Because of the higher basal ATPase rate in detergent, however, the stimulation effect is significantly less pronounced, corresponding to only a ∼50% increase for GSSG stimulation. Little change is observed for the K_m_s of MgATP between the presence and absence of glutathione derivatives for either detergent solubilized or nanodisc reconstituted *At*Atm3 (Figure 1).

**Figure 1.**
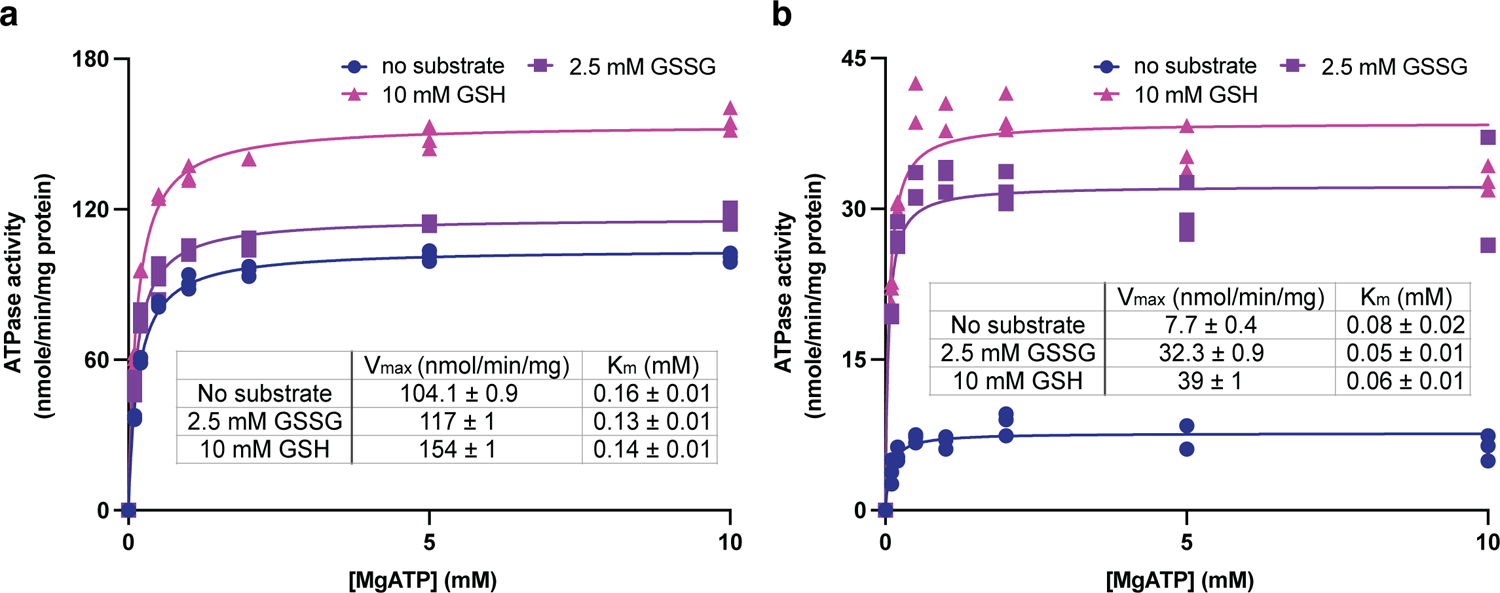
ATPase activities of *At*Atm3. ATPase activities measured in **a**) the detergent DDM and **b**) nanodiscs formed by membrane scaffolding proteins (MSP) and the lipid 1-palmitoyl-2-oleoyl-glyce- ro-3-phosphocholine (POPC). The ATPase activities were measured in the absence of substrate (●), at 2.5 mM GSSG (▪) and 10 mM GSH (▴). The corresponding values of Vmax and Km in different substrate conditions derived from fitting to the Michaelis-Menten equations are indicated. Each condition was measured three times with the individual data points displayed.

### Inward-facing, nucleotide-free conformational states

To map out the transport cycle, we attempted to capture *At*Atm3 in distinct liganded conformational states using single-particle cryoEM. We first determined the structure of *At*Atm3 reconstituted in nanodiscs at 3.4 Å resolution in the absence of either nucleotide or transport substrate (Figure 2a, S2). This structure revealed an inward-facing conformation for *At*Atm3 similar to those observed for the inward-facing conformations for *Sc*Atm1 (PDB ID: 4myc) and *Na*Atm1 (PDB ID: 6vqu) with overall alignment rmsds for the dimer of 2.6 Å (Figure S3a) and 2.1 Å (Figure S3b), respectively, and half-transporter alignment rmsds of 2.3 Å and 2.0 Å (Figure S3c), respectively. The primary distinction between these structures is the presence of a ∼20 amino acid loop between TM1 and TM2 of *At*Atm3 that would be positioned in the intermembrane space and is absent from the structures of *Sc*Atm1 and *Na*Atm1 (Figure 2a, S2de).

**Figure 2.**
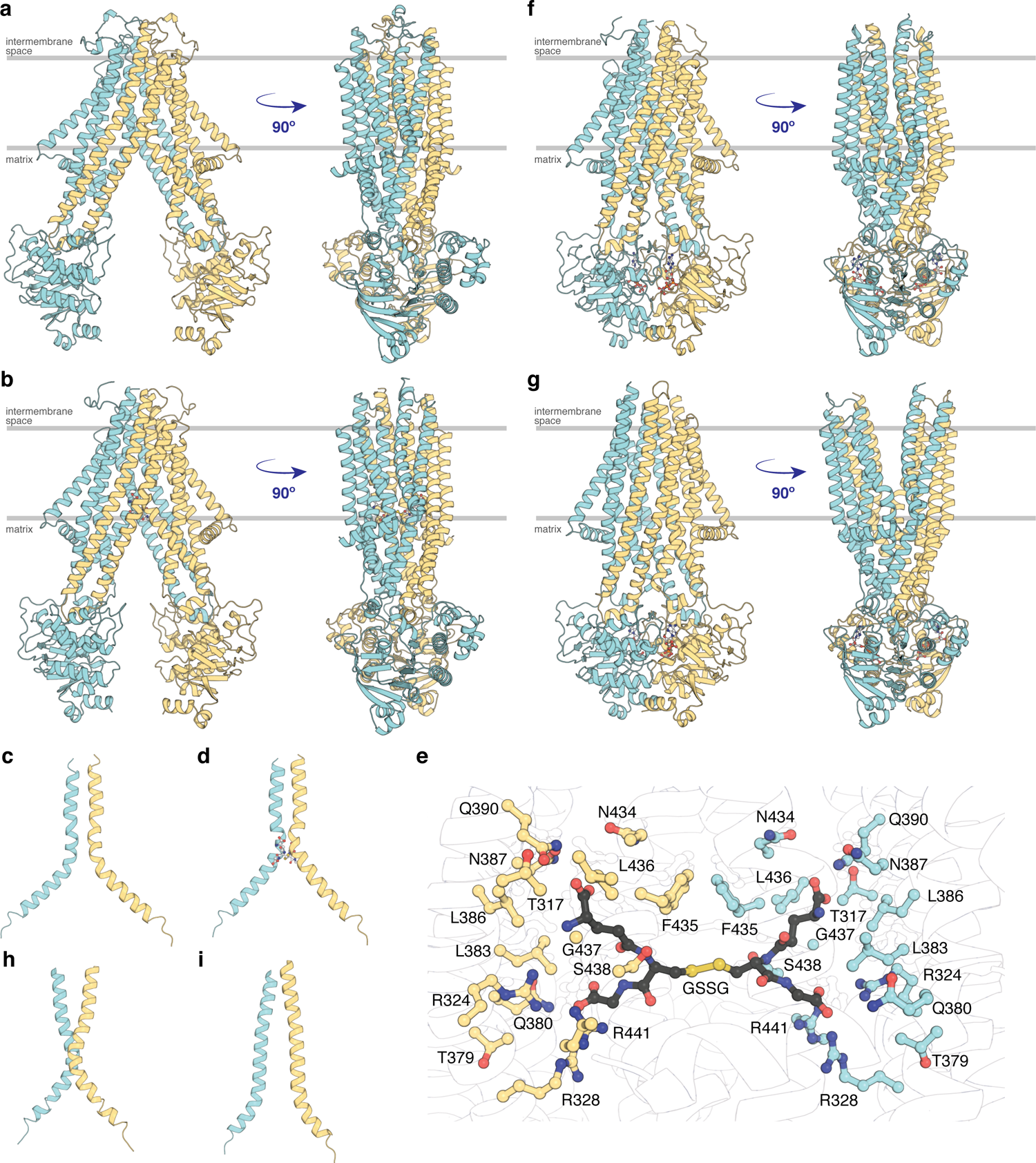
Structures of *At*Atm3. **a**) Inward-facing conformation in the apo state. **b**) Inward-facing conformation with GSSG bound. **c**) TM6s (residues 416-460) in the inward-facing conformation. **d**) TM6s in the GSSG-bound inward-facing conformation. The location of GSSG is indicated. **e**) Residues important in stabilizing GSSG binding site, identified by PDBePISA (Krissinel and Henrick, 2007). **f**) Closed conformation with MgADP-VO_4_ bound. **g**) Outward-facing conformation with MgADP-VO_4_ bound. **h**) TM6s in the closed conformation. **i**) TM6s in the outward-facing conformation.

To further look at the substrate binding, we determined a 3.6 Å resolution single-particle cryoEM structure of *At*Atm3 purified in DDM with bound GSSG (Figure 2b, S4). Although the overall resolution of the reconstruction was moderate (Figure S4d), we were able to model the GSSG molecule into the density map. In this structure, *At*Atm3 adopts an inward-facing conformation, with an overall alignment rmsd of 2.9 Å (Figure S5a) and half-transporter alignment rmsd of 1.6 Å (Figure S5b) comparing to the ligand-free inward-facing structure. The main difference between the two structures is the extent of NBD dimer separation (Figure S5a), where the GSSG bound structure presents a more closed NBD dimer relative to the substrate-free structure. As previously noted with *Na*Atm1 (Fan et al., 2020), the TM6 helices in these inward-facing structures of *At*Atm3 adopt a kinked conformation including residues 429-438 (Figure 2cd). This opens the backbone hydrogen bonding interactions to create the binding site for GSSG (Figure 2e) with binding pocket residues identified by PDBePISA (Krissinel and Henrick, 2007). The binding mode of GSSG in this *At*Atm3 inward-facing conformation is similar to that observed in the inward-facing structure of the GSSG bound *Na*Atm1 (Lee et al., 2014).

### MgADP-VO_4_ stabilized closed and outward-facing conformation

MgADP-VO_4_ has been found to be a potent inhibitor of multiple ATPases through formation of a stable species resembling an intermediate state during ATP hydrolysis (Crans et al., 2004; Davies and Hol, 2004). We determined two structures of *At*Atm3 stabilized with MgADP-VO_4_, one in the closed conformation with *At*Atm3 reconstituted in nanodiscs at 3.9 Å resolution (Figure 2f, S6), and the other in the outward-facing conformation with *At*Atm3 in DDM at 3.8 Å resolution (Figure 2g, S7). These two structures share an overall alignment rmsd of 1.7 Å with the primary difference being a change in separation of the TM helices surrounding the translocation pathway on the side of the transporter facing the intermembrane space (Figure S8). As a result of these changes in the TMDs, access to the intermembrane space is either blocked in the closed conformation (Figure 2f) or is open in the outward-facing conformation (Figure 2g). The changes in the TMDs are reflected in the conformations of TM6, which in the closed structure presents a kinked conformation (Figure 2h), in contrast to the straight conformation in the outward-facing structure that has the backbone hydrogen bonding interaction restored in the helices (Figure 2i). Further, the loops between TM1 and TM2 that are characteristic of the *At*Atm3 transporter are more ordered in the closed conformation than in the outward-facing conformation (Figure 2fg, S5-6). In contrast to the variation in the TMDs, the dimerized NBDs are virtually identical in these two structures with an overall alignment rmsd of 0.8 Å (Figure 2fg, S8).

## Discussion

The plant mitochondrial Atm3 transporter has been implicated in a diverse set of functions associated with transition metal homeostasis that are reflective of the roles that have been described for the broader Atm1 transporter family. To provide a general framework for addressing the detailed function of this transporter in plants, we have structurally and functionally characterized Atm3 from *Arabidopsis thaliana*. We first identified a construct of *At*Atm3 with the mitochondrial targeting sequence deleted that expressed well in *E. coli* (Figure S1). Following purification, the ATPase activities of *At*Atm3 were measured in both detergent and MSP nanodiscs as a function of MgATP concentrations (Figure 1). Overall, the ATPase rate measured in detergent is about 5-fold greater than that measured in nanodiscs, perhaps indicative of a more tightly coupled ATPase activity in a membrane-like environment. Both GSH and GSSG stimulate the ATPase activity by increasing V_max_, with little change observed in the K_m_ for MgATP. ABC transporters are typically envisioned as utilizing an ‘alternating access’ mechanism, in which the substrate-binding site transitions between inward- and outward-facing conformations coupled to the binding and hydrolysis of ATP. In an idealized two-state model, ABC transporters only adopt these two limiting conformations, but structural characterizations of ABC transporters in the presence of nucleotides and substrate analogs have identified a variety of intermediates, including occluded (with a ligand binding cavity exhibiting little or no access to either side of the membrane) and closed (no ligand binding cavity) conformations. The most extensive analysis of the conformational states of an Atm1 type exporter has been detailed for *Na*Atm1 and assigned to various states in the transport cycle (Fan et al., 2020; Lee et al., 2014). In the present work, we have determined four structures of *At*Atm3 in three different conformational states by single particle cryo-EM: two inward-facing conformations (with and without bound GSSG) (Figure 2ab), together with closed and outward-facing states stabilized by MgADP-VO_4_ (Figure 2fg). The parallels between the structurally characterized conformations of *At*Atm3 and *Na*Atm1 support the idea that these conformational states are relevant to the transport cycle, and not simply an artifact of the specific conditions used to prepare each sample. The conformations observed for *At*Atm3 and *Na*Atm1 do not completely correspond, however; most notably, the outward-facing conformation observed for *At*Atm3 had not been previously observed with *Na*Atm1 (Fan et al., 2020; Lee et al., 2014), while the occluded conformations found with *Na*Atm1 were not observed for *At*Atm3. The major differences between the closed and outward-facing conformations of *At*Atm3 stabilized with MgADP-VO_4_ are in the arrangements of the TMDs, while the NBDs are closely superimposable. In contrast, the primary differences between the two structures of inward-facing conformations of *At*Atm3 are in the relative positioning of the NBDs which are more widely separated in the apo structure relative to the GSSG bound structure; the arrangements of the TMDs in the dimer are similar in both structures.

The conformational changes in the TMDs underlying the transport cycle are associated with changes in the extent of kinking of TM6 and the positioning of TM4-TM5 relative to the core formed by the remaining four TM helices. As noted for *Na*Atm1, we observed kinked TM6s in the inward-facing and closed state of *At*Atm3 (Figure 2cdh), but not the outward-facing conformation (Figure 2i). These conformational changes lead to changes in the volume of the central cavity forming the glutathione binding site. Using the program CastP (Tian et al., 2018) with a probe radius of 2.5 Å, the cavity volumes of the inward-facing apo and GSSG bound structures were measured to be ∼6,500 Å^3^ (Figure 3a) and ∼4,300 Å^3^ (Figure 3b), respectively, while the closed conformation exhibits a cavity volume of ∼300 Å^3^ (Figure 3c), and the outward-facing conformation has a cavity volume of ∼5,700 Å^3^ (Figure 3d). We also measured the accessible solvent areas (ASA) of the key residues forming the binding site for GSSG in the different conformational states using Areaimol in CCP4 (Winn et al., 2011); the ASA of the inward-facing, inward-facing with GSSG bound, closed and outward-facing structures are ∼1,500 Å^2^, ∼1,100 Å^2^, ∼900 Å^2^, and ∼1,300 Å^2^, which are also highly correlated with the cavity volume calculations. Most of the binding pocket residues remain exposed in all conformations with a few having large relative changes than others (Figure S9). Further, the cavity volume measurements are comparable to those calculated for *Na*Atm1 (Fan et al., 2020). The similarities in conformational states between *Na*Atm1 and *At*Atm3 indicate these transporters follow the same basic mechanism, in which straightening of TM6s in the transition from inward- to outward-conformation leads to the release of substrate to the opposite side of the membrane. Following substrate release, the transporter resets to the inward-facing conformation through the closed conformation adopted after ATP hydrolysis; the decreased size of the substrate binding cavity helps enforce substrate release and unidirectionality of substrate transport.

**Figure 3.**
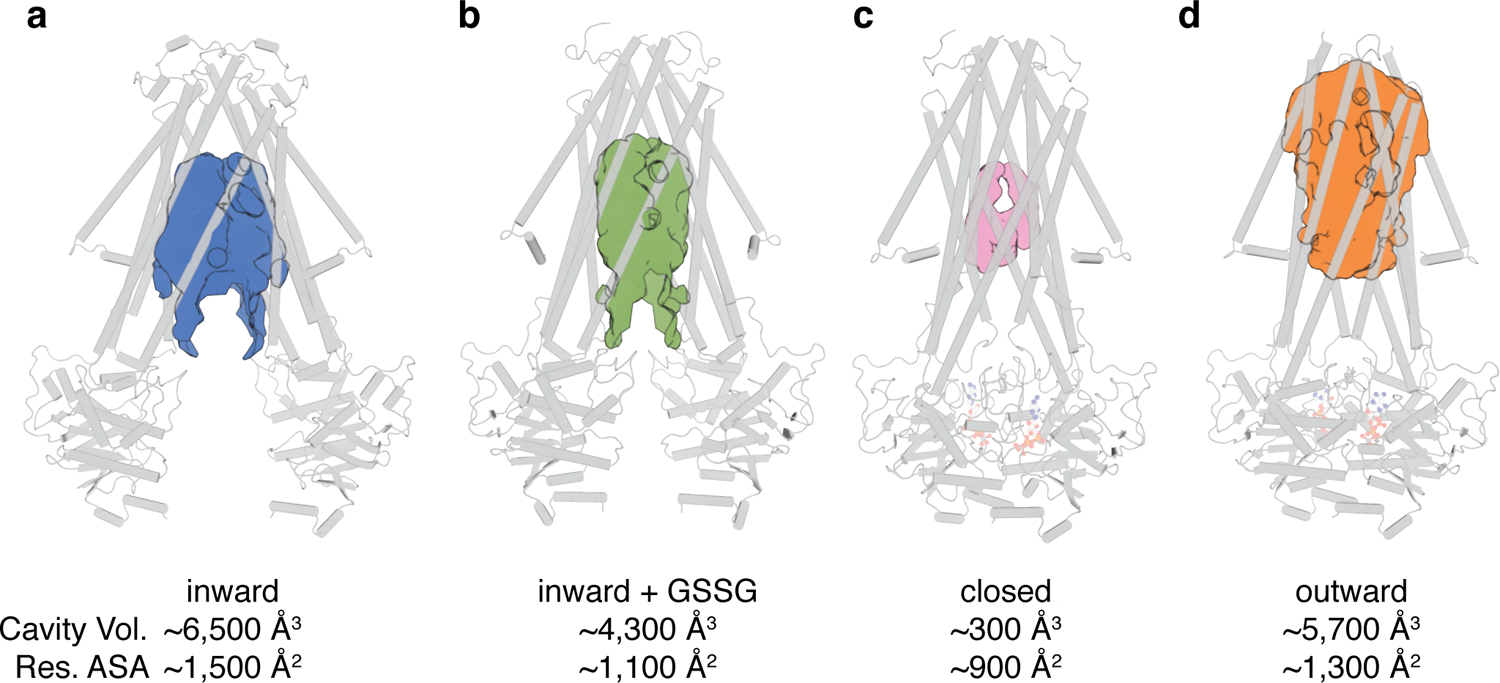
Binding cavity analysis. **a**) Central cavity of the apo inward-facing conformation. **b**) Central cavity of the inward-facing conformation with GSSG bound. **c**) Closed conformation with no central cavity observed. **d**) Central cavity of the outward-facing conformation. Cavity volumes were measured by CastP (Tian et al., 2018) using a probe radius of 2.5 Å. *At*Atm3 is shown as a grey cartoon representation, while cavities are depicted as color surfaces. The accessible solvent areas (ASA) of the key residues in the GSSG binding pockets of different structures were calculated Areaimol in CCP4 (Winn et al., 2011).

The binding pocket for GSSG identified in this work primarily consists of residues from TM5 and TM6, with additional contributions from residues in TM3 and TM4. The GSSG binding site for *At*Atm3 largely overlaps with that identified previously for *Na*Atm1 (Lee et al., 2014) and for the binding of reduced GSH to *Sc*Atm1 (Srinivasan et al., 2014). Inspection of a sequence alignment of Atm1 homologs (Figure S10) reveals that those residues forming the glutathione binding site are largely conserved, particularly if they are involved in polar interactions. A striking feature is the stretch of residues from P432 to R441 in the middle of TM6 (*At*Atm3 sequence numbering) with sequence PLNFLGSVYR with a high degree of sequence conservation. P432 is associated with the TM6 kink in inward-facing conformations that permits formation of hydrogen bonds between exposed peptide groups with GSSG (Lee et al., 2014); as TM6 straightens in the occluded and outward-facing conformations, these peptide groups are no longer available to bind the transport substrate (Fan et al., 2020). A sequence alignment of the structurally characterized *At*Atm3, *Na*Atm1, *Sc*Atm1 and human ABCB7 and ABCB6 (Figure S2) transporters establishes that residues in the binding pockets are conserved, including T317, R324, R328, N387, Q390, L433, G437 and R441 (Figure S10). The conservation of binding pocket residues as calculated by the program ConSurf (Landau et al., 2005) is illustrated in Figure 4 suggests that the substrates for these transporters may share common features, such as the glutathione backbone. Positions such where sequence variability is evident, such as residue 435 (Figure 4B and S9), may reflect the binding of distinct GSSG-derivatives by different eukaryotic and prokaryotic homologs.

**Figure 4.**
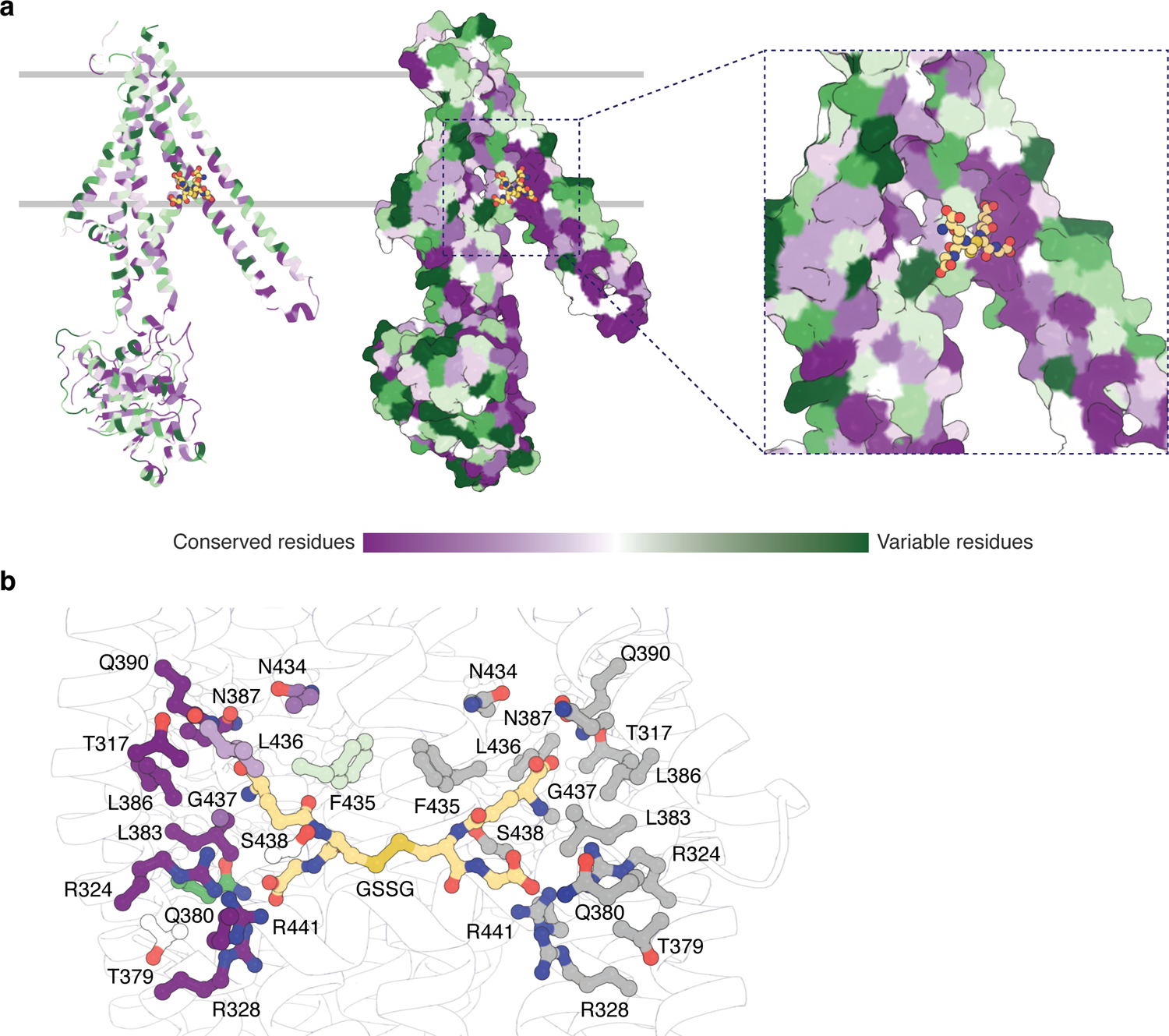
Substrate binding site conservation. **a**) Sequence conservation of *At*Atm3, *Na*Atm1, *Sc*Atm1, human ABCB7 and human ABCB6 calculated by ConSurf (Landau et al., 2005) plotted on a cartoon and surface representations of a half-transporter of *At*Atm3 in the GSSG bound inward-facing conformation. GSSG is shown in spheres. **b**) Conservation of key residues in the GSSG binding pocket. Residues in one chain is colored based on the conservation and residues in the second chain is colored in grey. All residues and GSSG are shown in ball and sticks.

An important property of disulfide containing compounds such as oxidized glutathione is that they can undergo disulfide – thiol exchange with free -SH groups (Creighton, 1984; Nagy, 2013). This reactivity creates potential challenges for proteins such as *At*Atm3 since reaction of a disulfide containing ligand such as GSSG with the thiol-containing side chain of cysteine could lead to formation of the mixed disulfide, thereby covalently connecting glutathione to the protein and releasing reduced GSH. Formation of the covalently linked mixed disulfide would be expected to restrict the access of exogenous ligands to the substrate binding cavity and hence would inhibit transport. For membrane proteins, cysteine residues are present in transmembrane helices with a frequency of about 1% (Baeza-Delgado et al., 2013). Although no cysteines residues are present in the *At*Atm3 binding pocket, we analyzed additional Atm3 homologs from plants. For this analysis, we used the NCBI blastp server (Altschul et al., 1997) and selected 410 sequences with a sequence identity of 50% to 100% and query coverage of 80% to 100% with *At*Atm3, Within the six TM helices, the overall presence of cysteines was found to be ∼0.4%. In this alignment, no cysteines were found in residues forming the glutathione binding cavity (Figure 5); more strikingly, no cysteine residues were found at any position of TM6 for these homologs (Figure 5f). Cysteines that are present in the TMs are either distant from the binding site, such as position 405 in TM5 of *At*Atm3 (Figure 5eg) or if they are closer to the binding site, are positioned on the opposite side of the TMs, such as positions 149, 215, 290 and 307 (Figure 5abcd). The lack of cysteines (Figure 5g) may reflect a design feature in the glutathione binding site, namely cysteines that could potentially form inhibitory mixed disulfides during the transport cycle are excluded from Atm3.

**Figure 5.**
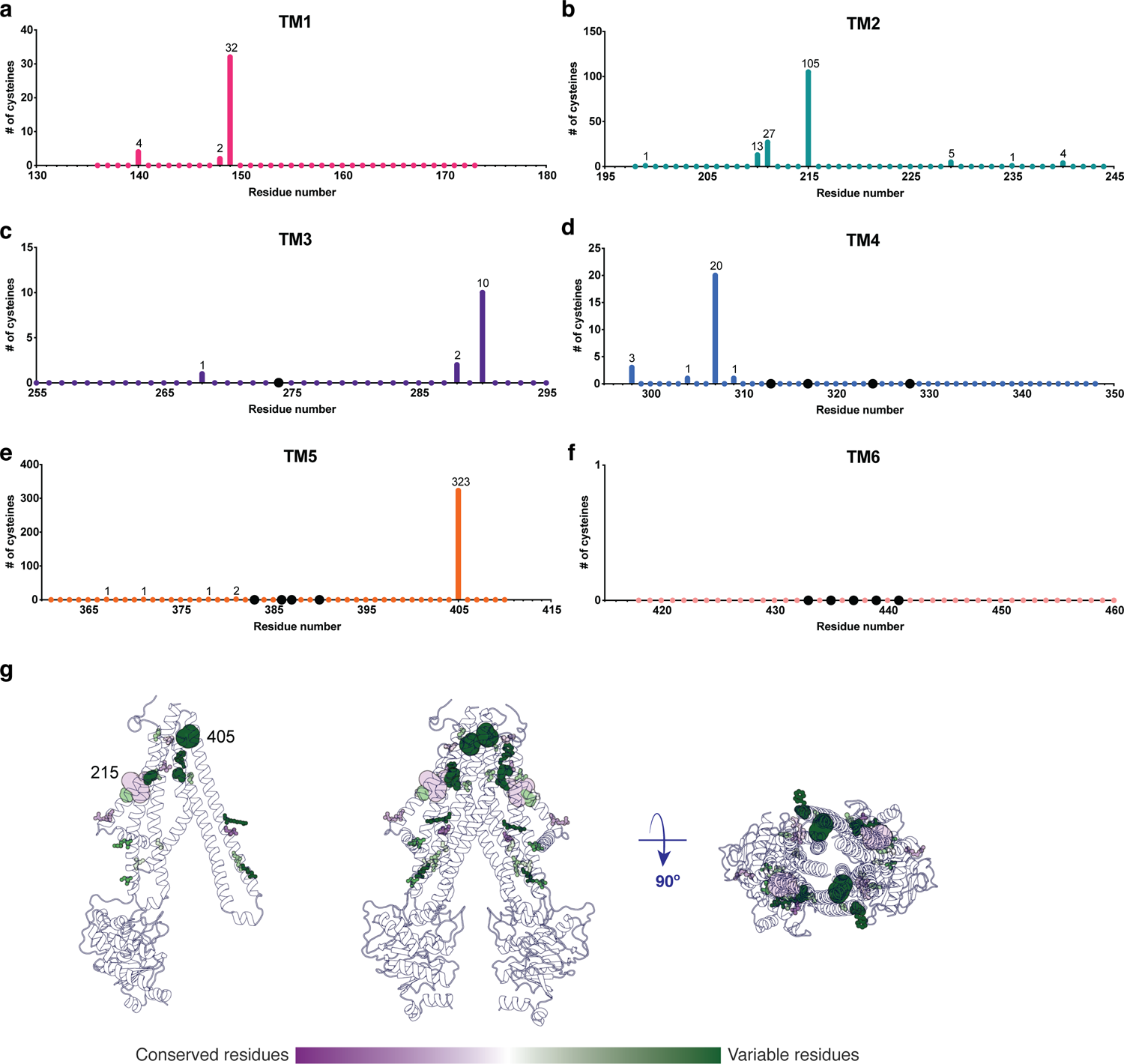
Cysteine residues found in transmembrane helices in an alignment of 410 *At*Atm3 homologs. **a-f**) Residues are numbered based on the *At*Atm3 sequence. Small colored dots represent different residue positions, while the larger black dots indicate residues observed to interact with glutathione in the *Na*Atm1 and *At*Atm3 structures. The numbers above a given residue indicate the number of sequences in the alignment with a cysteine at that position; unlabeled positions denote positions where no cysteines were observed in the alignment. **g**) Transmembrane cysteine residues in the inward-facing conformation of *At*Atm3. All positions with cysteine counts are shown in spheres. Positions that have 1 to 10 counts of cysteines are shown in small spheres, positions that have 11 to 100 counts of cysteins are shown in medium sized spheres, and positions with 101 and above counts of cysteines are shown in large spheres. The spheres are colored based on the consurf coloring in Figure 4. The distance between the Cα’s of two 215 residues is 37 Å, and the distance between the Cα’s of two 405 residues is 25 Å.

As a general strategy to stabilize ABC transporters in distinct conformational states, different nucleotides or transport substrates are mixed with the transporter. The expectation is that a particular set of ligands will stabilize a specific conformational state, and so we were surprised to have captured with MgADP-VO_4_ both a closed conformation in MSP nanodiscs and an outward-facing conformation in detergent. Given the similarities in the NBDs between these two structures, the distinctive arrangements of the TMDs between the closed and outward-facing conformations presumably arises from differences in the TMD environment provided by MSP nanodiscs and DDM, respectively. Furthermore, this phenomenon of obtaining different conformational states with MgADP-VO_4_ is not unprecedented, however, since previously determined structures of MgADP-VO_4_ stabilized ABC exporters include *Na*Atm1 in the closed conformation (Fan et al., 2020), *Thermus thermophilus* TmrAB in the occluded and outward-facing conformations (Hofmann et al., 2019) and *Escherichia coli* MsbA in the closed conformation (Mi et al., 2017).

Despite extensive efforts, we were unable to prepare the ternary complex of *At*Amt3 with bound GSSG and MgATP. To assess more generally the relationship between the conformational states of ABC exporters and the presence or absence of nucleotide and transport substrate, we systematically compared the conformations of 80 half-transporters from the available structures of type IV ABC transporters (Table S2). To define the conformational state in a consistent fashion, we performed a principal component analysis (PCA) of these half-transporter structures (see Material and Methods); the principal component is dominated by the conformational state of the TMDs, which represents ∼62% of total conformational change from the inward-facing to the outward-facing conformation. The distribution of component 1 of the PCA is shown in Figure 6 with the most extreme inward-facing conformations to the left and the outward-facing conformations to the right.

**Figure 6.**
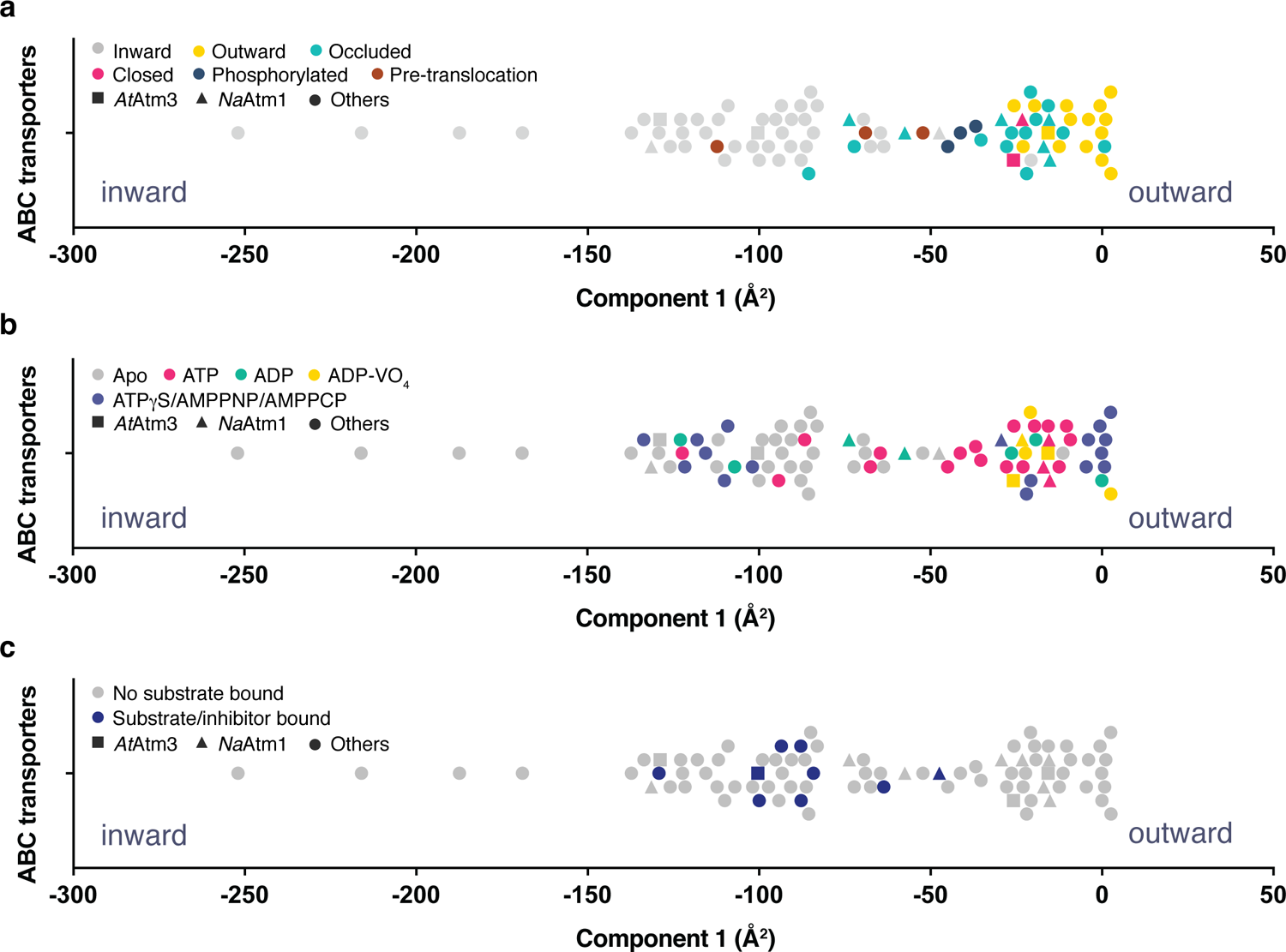
Principal component analysis (PCA) of type I ABC exporters. **a)** 1-dimensional plot of component 1, colored according to the conformational states assigned in the original publications. The plot is oriented with the most inward and most outward conformations to the left and right, respectively. **b)** 1-dimensional plot of component 1 with inward-facing to outward-facing structures, colored by their nucleotide states. **c)** 1-dimensional plot of component 1 with inward-facing to outward-facing structures, colored by their substrate states. Each marker (triangle (▲), square (▪), circle (●)) represents a unique half-transporter structure. Triangles (▲) represent structures of *At*Atm3, squares (▪) represent structures of *Na*Atm1 and circles (●) represent structures for other ABC transporters.

To validate this approach, we color coded each structure according to their published conformational state (Figure 6a). It is evident that outward-facing conformations occur on the right-handed side of the figure with component 1 values between −25 and 0 Å^2^, while occluded, closed and inward-facing conformations cluster around −25, −25 and −100 Å^2^, respectively. Hence, the magnitude of the principal component does capture the expected trends in conformational state, with increasing values corresponding to a progression from inward-facing to outward-facing conformations. The correspondence is not exact, however, since assigned conformational states overlap, which could reflect either limitations of the principal component analysis, and/or inconsistent assignments of conformational states between different structures. We further note that some conformational states have a very wide distribution, particularly the inward-facing conformations, with several structures exhibiting widely separated subunits (with values for component 1 below −150 Å^2^). Using the principal component analysis, we could relate protein conformation to binding of nucleotides (Figure 6b) and substrates/inhibitors (Figure 6c). This analysis establishes that occluded and outward-facing conformations are largely nucleotide bound with either ATP or an ATP analog, but there are exceptions, most notably the original ADP bound structure of Sav1866 ((Dawson and Locher, 2006); a similar Sav1866 structure was subsequently solved with bound AMPPNP (Dawson and Locher, 2007)). Inward-facing conformations are observed in both nucleotide-free and nucleotide bound forms (Figure 6b). Only the MgADP-VO_4_ bound structures are exclusively found to occupy a small conformational space in the closed/outward-facing region. In contrast to the binding of nucleotides to all conformational states of these transporters, a distinct pattern is observed for the binding of transport substrates where the transporter invariably adopts the inward-facing conformation (Figure 6c). Intriguingly, no structures to date have been published that contain both nucleotide and transport substrate, suggesting that this ternary complex exists only transiently during the transport cycle. As this is a key intermediate for understanding how the binding of transport substrate stimulates the ATPase activity, characterization of the ternary complex represents an outstanding gap in the mechanistic characterization of ABC exporters.

## Material and Methods

### Cloning, expression and purification

A pET-21a (+) plasmid containing the full-length *Arabidopsis thaliana* Atm3 (*At*Atm3) gene with a C-terminal 6x-His tag was purchased from Genscript (Genscript, NJ). Mutagenesis reactions generating the N-terminal 60, 70 and 80 amino acids deletion mutants were carried out with the Q5 mutagenesis kit (New England Biolabs, MA). All *At*Atm3 constructs were overexpressed in *Escherichia coli* BL21-gold (DE3) cells (Agilent Technologies, CA) using ZYM-5052 autoinduction media as described previously (Fan et al., 2020). Cells were harvested by centrifugation and stored at −80 °C.

For purification, frozen cell pellets were resuspended in lysis buffer containing 100 mM NaCl, 20 mM Tris, pH 7.5, 40 mM imidazole, pH 7.5, 10 mM MgCl_2_ and 5 mM b-mercaptoethanol (BME) in the presence of lysozyme, DNase, and cOmplete protease inhibitor tablet (Roche, Basel, Switzerland). The resuspended cells were lysed with an M-110L pneumatic microfluidizer (Microfluidics, MA). Unlysed cells and cell debris were removed by centrifugation at ∼20,000 x *g* for 30 minutes at 4 °C. The membrane fraction containing *At*Atm3 was collected by ultracentrifugation at ∼113,000 x *g* for an hour at 4 °C. The membrane fraction was then resuspended in buffer containing 100 mM NaCl, 20 mM Tris, pH 7.5, 40 mM imidazole, pH 7.5, and 5 mM BME and further solubilized by stirring with the addition of 1% n-dodecyl-β-D-maltopyranoside (DDM) (Anatrace, OH) for an hour at 4 °C. The DDM solubilized membrane was ultracentrifuged at ∼113,000 x *g* for 45 minutes at 4 °C to remove any unsolubilized material. The supernatant was loaded onto a prewashed NiNTA column. NiNTA wash buffer contained 100 mM NaCl, 20 mM Tris, pH 7.5, 50 mM imidazole, pH 7.5, 5 mM BME and 0.02% DDM, while the elution buffer contained the same components, but with 380 mM imidazole. The eluent was subjected to size exclusion chromatography using HiLoad 16/60 Superdex 200 (GE Healthcare, IL) with buffer containing 100 mM NaCl, 20 mM Tris, pH 7.5, 5 mM BME and 0.02% DDM. Peak fractions were collected and concentrated to ∼10 mg/ml using Amicon concentrators (Millipore, MA).

### ATPase activity assay

ATPase assays were carried out as described previously for both the detergent purified and the reconstituted nanodisc samples (Fan et al., 2020) using a molybdate based colorimetric assay (Chifflet, 1988).

### Nanodisc reconstitution

For the *At*Atm3 structures in nanodiscs, the reconstitution was performed following the previously described protocol (Fan et al., 2020). The reconstitution was done with a 1:4:130 molar ratio of *At*Atm3: membrane scaffolding protein (MSP) 1D1: 1-palmitoyl-2-oleoyl-glycero-3-phosphocholine (POPC) (Avanti Polar Lipids, AL). After overnight incubation at 4 °C, the samples were subjected to size exclusion chromatography with a Superdex 200 Increase 10/300 column (GE Healthcare, IL). The peak fractions were directly used for grid preparation with the reconstituted samples at ∼0.5 mg/ml. For the structure of *At*Atm3 with GSSG bound in the inward-facing conformation, the detergent purified protein was incubated with 10 mM GSSG, pH 7.5 at 4 °C for an hour with *At*Atm3 at 4 mg/ml before freezing grids.

### Grid preparation

For the *At*Atm3 structure with MgADP-VO_4_ bound in the outward-facing conformation, the detergent purified protein was incubated with 4 mM ATP, pH 7.5, 4 mM MgCl_2_ and 4 mM VO_4_^3-^ with protein at 5 mg/ml at 4 °C overnight before freezing grids. For all grids, 3 μL of protein solution was applied to freshly glow-discharged UltrAuFoil 2/2 200 mesh grids (apo inward-facing conformation and closed conformation, both in nanodiscs) and UltrAuFoil 1.2/1.3 300 mesh grids (Electron Microscopy Sciences, PA) (GSSG bound inward-facing conformation and outward-facing conformation, both in detergent) and blotted for 4 to 5 seconds with a blot force of 0 and 100% humidity at room temperature using the VitroBot Mark IV (Thermo-Fisher, MA).

### Single-particle cryoEM data collection, processing and refinement

Datasets for the inward-facing conformations in apo and GSSG bound states, and the outward-facing conformation with MgADP-VO_4_ bound were collected with a Gatan K3 direct electron detector (Gatan, CA) on a 300 keV Titan Krios (Thermo-Fisher, MA) in the super-resolution mode using SerialEM at the Caltech CryoEM facility. These datasets were collected using a defocus range between −1.5 to −3.0 μm and a total dosage of 60 e^-^/Å^2^. The dataset for the closed conformation with MgADP-VO_4_ bound was collected with a Falcon 4 direction electron detector (Thermo-Fisher, MA) on a 300 keV Titan Krios (Thermo-Fisher, MA) in the super-resolution mode using EPU (Thermo-Fisher, MA) at the Stanford-SLAC Cryo-EM Center (S2C2) with a defocus range between −1.5 and −2.1 μm and a total dosage of ∼48 e^-^/Å^2.^

Detailed processing workflows of all single-particle cryoEM datasets are included in Figure S2, S4, S6 and S7. Datasets for the inward-facing conformation in apo and GSSG bound states, and the outward-facing conformation with MgADP-VO_4_ bound were motion corrected with the patch motion correction in cryoSPARC 2 (Punjani et al., 2017), while the dataset for the closed conformation with MgADP-VO_4_ bound was motion corrected with motioncor2 (Zheng et al., 2017). The subsequent processing of all datasets was performed in a similar fashion. The contrast transfer function (CTF) parameters were estimated with patch CTF estimation in cryoSPARC 2 (Punjani et al., 2017). Particles were picked with blob picker using a particle diameter of 80 to 160 Å and then extracted. Rounds of two-dimensional (2D) and three-dimensional (3D) classifications were performed, leaving 157,762, 259,020, 140,569 and 103,161 particles for the inward-facing apo, inward-facing GSSG bound, closed, and outward-facing conformations, respectively. The final reconstructions were refined with homogeneous, non-uniform and local refinements in cryoSPARC 2 with C2 symmetry (Punjani et al., 2017). The masks used in local refinements were generated in Chimera (Pettersen et al., 2004).

The initial model of the *At*Atm3 inward-facing conformation in apo state was obtained using the inward-facing occluded structure of *Na*Atm1 (PDB ID: 6pam) as a starting model (Fan et al., 2020). The model fitting was carried out with phenix.dock_in_map (Liebschner et al., 2019). That apo inward-facing conformation model of *At*Atm3 was subsequently used as the starting model for the inward-facing GSSG bound structure. The previous *Na*Atm1 closed conformation structure (PDB: 6par) was used as the starting model for both the closed and the outward-facing conformations. Model building and ligand fitting were carried out manually in *Coot* (Emsley et al., 2010) and the structures were refined with phenix.real_space_refine (Liebschner et al., 2019).

### Structure superposition

Structure superpositions for calculating the root mean square deviations (rmsds) between different structures were performed with the SSM option in *Coot* (Emsley et al., 2010).

### Principal component analysis

The objective of our principal component analysis (PCA) was to provide a quantitative foundation for relating the binding of nucleotides and transported ligands to the conformational state of type IV ABC transporters that include *At*Atm3 and related Atm1 transporters (Thomas et al., 2020). For this purpose, we first identified 10 polypeptide stretches containing 7 to 21 residues at equivalent positions in 80 structurally characterized type IV transporters (Table S2), including residues from the TMD and from the NBD. In this sequence selection process, a single half-transporter was used for homodimeric transporters, and both half-transporters were used for heterodimeric transporters, whether encoded by two different half-transporter peptides or on a single peptide. The coordinates of C*α* positions for the selected residues were extracted and aligned to that of the outward-facing conformation of Sav1866 (PDB ID: 2hyd) based on the C*α* coordinates in TM3 and TM6, which were previously found to provide a useful reference frame for studying conformational changes (Lee et al., 2014). *At*Atm3 residues used in alignment: 140-160, 225-245, 255-275, 322-342, 362-382, 423-443, 504-513, 517-524, 618-632 and 681-688. *Na*Atm1 residues used in alignment: 36-56, 107-127, 137-157, 204-224, 244-264, 305-325, 386-395, 399-406, 500-514 and 563-570.

The PCA was performed using the “essential dynamics” algorithm (Amadei et al., 1993). The full transporter was used in these calculations with the outward-facing state of Sav1866 (PDB ID: 2hyd) serving as the reference state. The first component captured 62% of the overall conformational variation among these structures, and so the eigenvalues corresponding to this component were used to order the different structures along a single axis (Figure 6). In general, the conformational states assigned to each structure parallel those obtained from the PCA analysis; differences likely reflect the absence of standardized definitions for assigning the conformational states of ABC transporters as well as the limitations of this PCA analysis, particularly the use of only the dominant eigenvector.

## Author contributions

C.F. and D.C.R. designed the research; C.F. performed the research; C.F. and D.C.R. analyzed the data; and C.F and D.C.R. prepared the manuscript.

## Data availability

The atomic coordinates for inward-facing, inward-facing with GSSG bound, closed and outward-facing conformations were separately deposited in the Protein Data Bank (PDB) and the Electron Microscopy Data Bank (EMDB) with accession codes: PDB 7N58, 7N59, 7N5A and 7N5B; EMDB EMD-24182, EMD-24183, EMD-24184 and EMD-24185. The plasmid encoding full-length *At*Atm3 and the *At*Atm3 with N-terminal 80 residue deletion were deposited in Addgene with Addgene ID 172321 and 173045, respectively. The raw data for ATPase assays presented in Figure 1 are provided in Supplementary File 1, while the essdyn.f Fortran source code used for the PCA analysis is provided as Source Code 1.

## Acknowledgments

We thank Andrey Malyutin, Songye Chen and Corey Hecksel for their support during single-particle cryo-EM data collections. Cryo-electron microscopy was performed in the Beckman Institute Resource Center for cryo-Electron Microscopy at Caltech and at the Stanford-SLAC Cryo-EM Center (S2C2). The S2C2 is supported by the National Institutes of Health Common Fund Transformative High Resolution Cryo-Electron Microscopy program. We thank the Beckman Institute for their support of the cryo-EM facility at Caltech.

---

**Figure S1.**
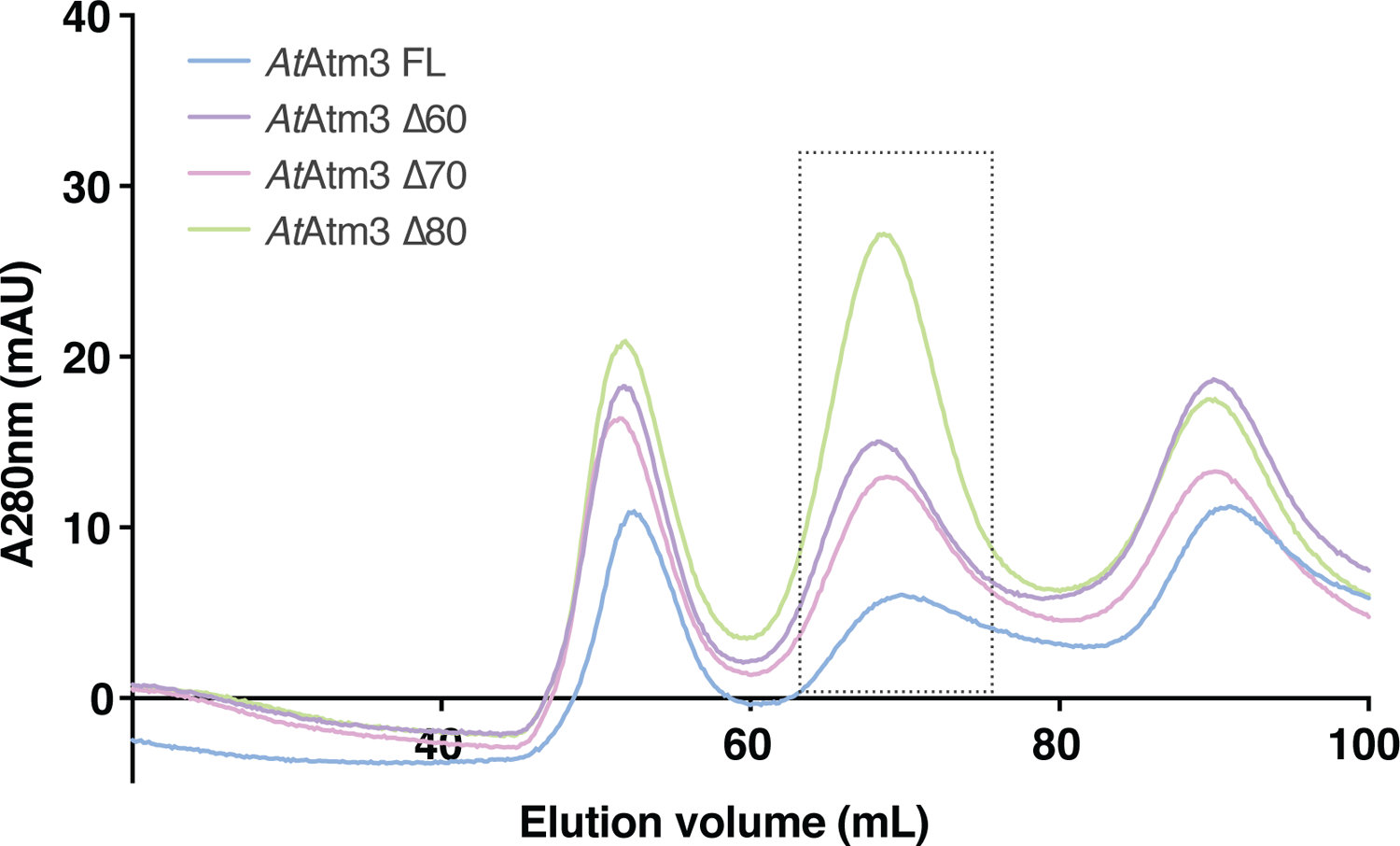
*At*Atm3 constructs. Size exclusion chromatography profile of *At*Atm3 constructs using HiLoad 16/60 Superdex 200 (GE Healthcare). Fractions collected for structural and fuctional analysis are boxed with dotted line.

**Figure S2.**
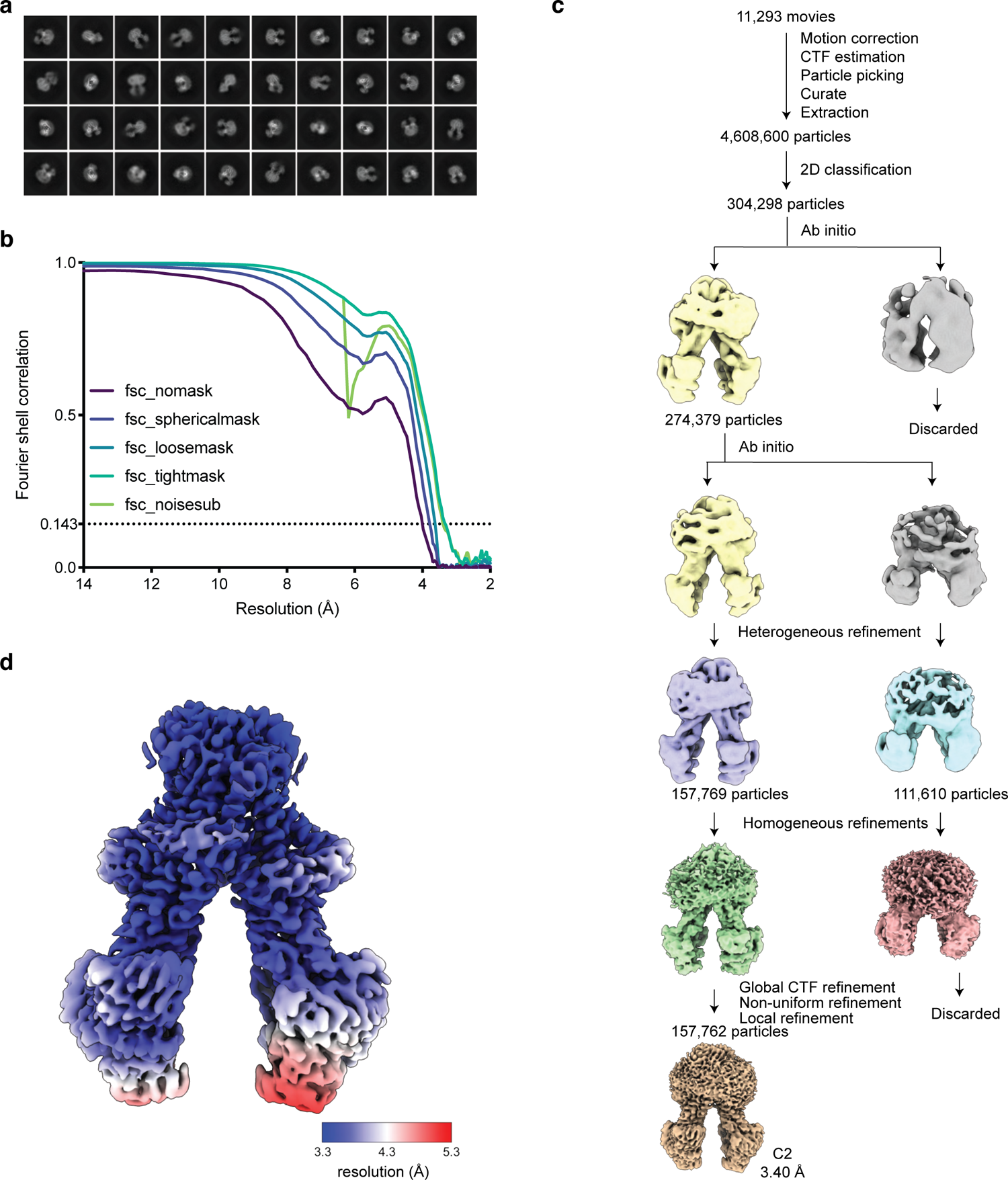
Single particle cryoEM structure of *At*Atm3 in the inward-facing conformation. **a**) Examples of 2D classes, **b**) FSC curves showing the resolution estimate for the final reconstruction, **c**) workflow of single-particle data processing, and **d**) local resolution estimation of the *At*Atm3 inward-facing conformation.

**Figure S3.**
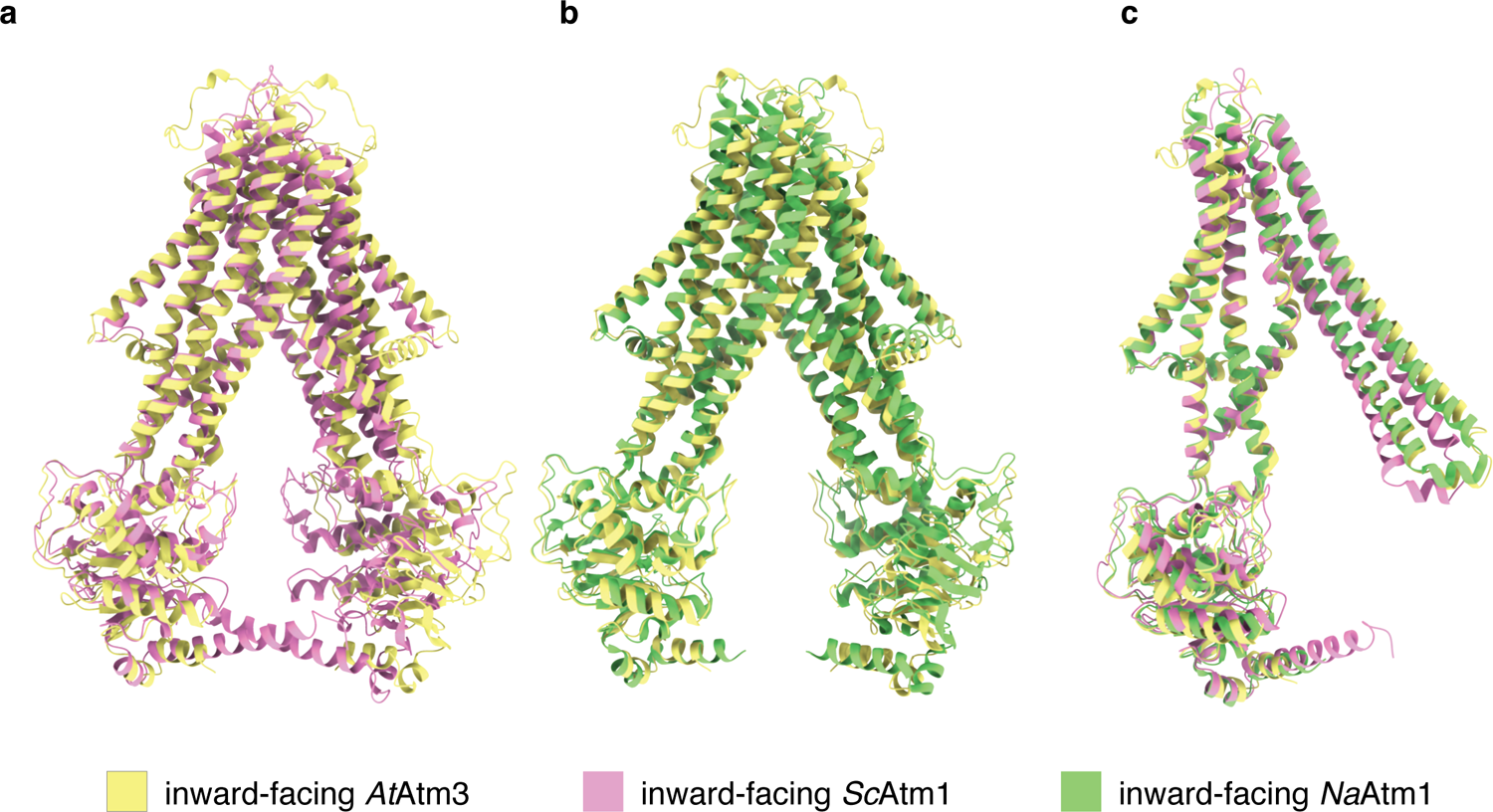
Structural alignment of *At*Atm3 to other ATM transporters. **a**) Overall alignment of inward-facing *At*Atm3 to *Sc*Atm1 (PDB ID: 4myc) with an overall RMSD of 2.6 Å. **b**) Overall alignment of inward-facing *At*Atm3 to *Na*Atm1 (PDB ID: 6vqu) with an overall RMSD of 2.1 Å. **c**) Half-transporter alignments *Sc*Atm1 (PDB ID: 4MYC) and *Na*Atm1 (PDB ID: 6vqu) to inward-facing *At*Atm3 to with RMSDs of 2.3 Å and 2.0 Å, separately.

**Figure S4.**
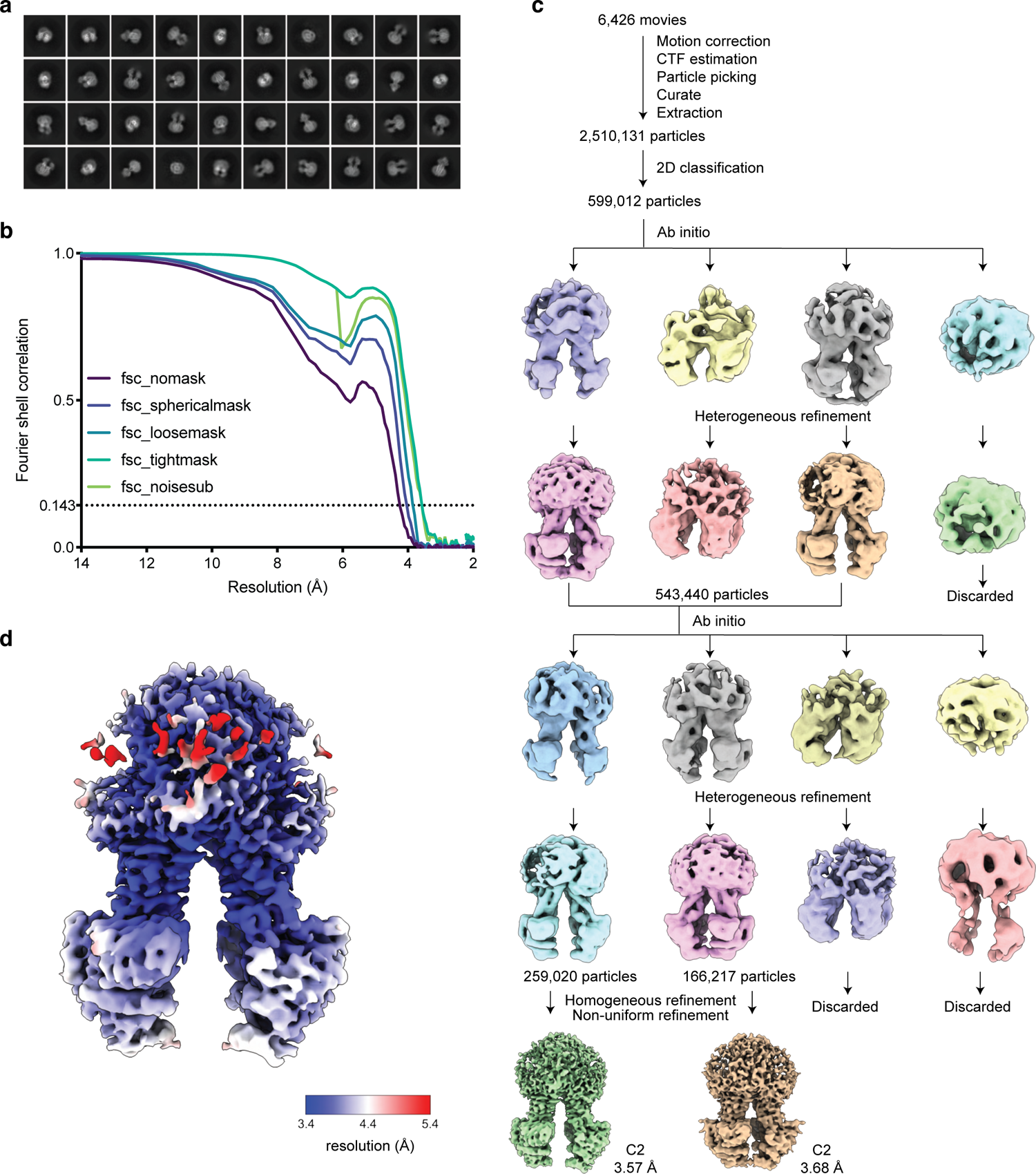
Single particle cryoEM structure of *At*Atm3 in the inward-facing conformation with GSSG bound. **a**) Examples of 2D classes, **b**) FSC curves showing the resolution estimate for the final reconstruction, **c**) workflow of single-particle data processing, and **d**) local resolution estimation of the *At*Atm3 inward-facing conformation with GSSG bound.

**Figure S5.**
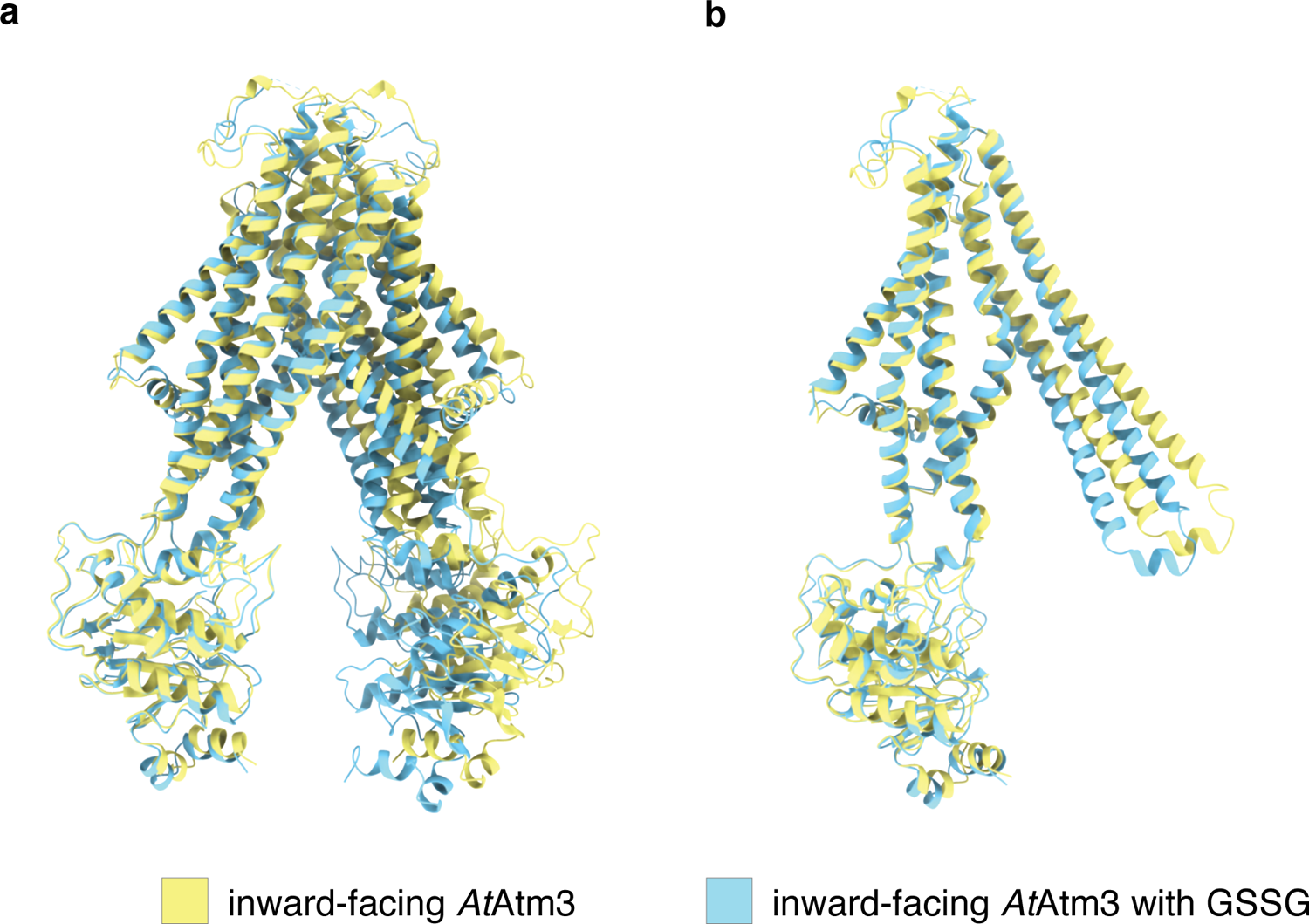
Structural alignment of *At*Atm3 in the inward-facing conformation. **a**) Overall alignment of inward-facing *At*Atm3 in the apo state to the GSSG bound state with an overall RMSD of 2.9 Å. **b**) Half-transporter alignment inward-facing *At*Atm3 in the apo state to the GSSG bound state with an RMSD of 1.6 Å.

**Figure S6.**
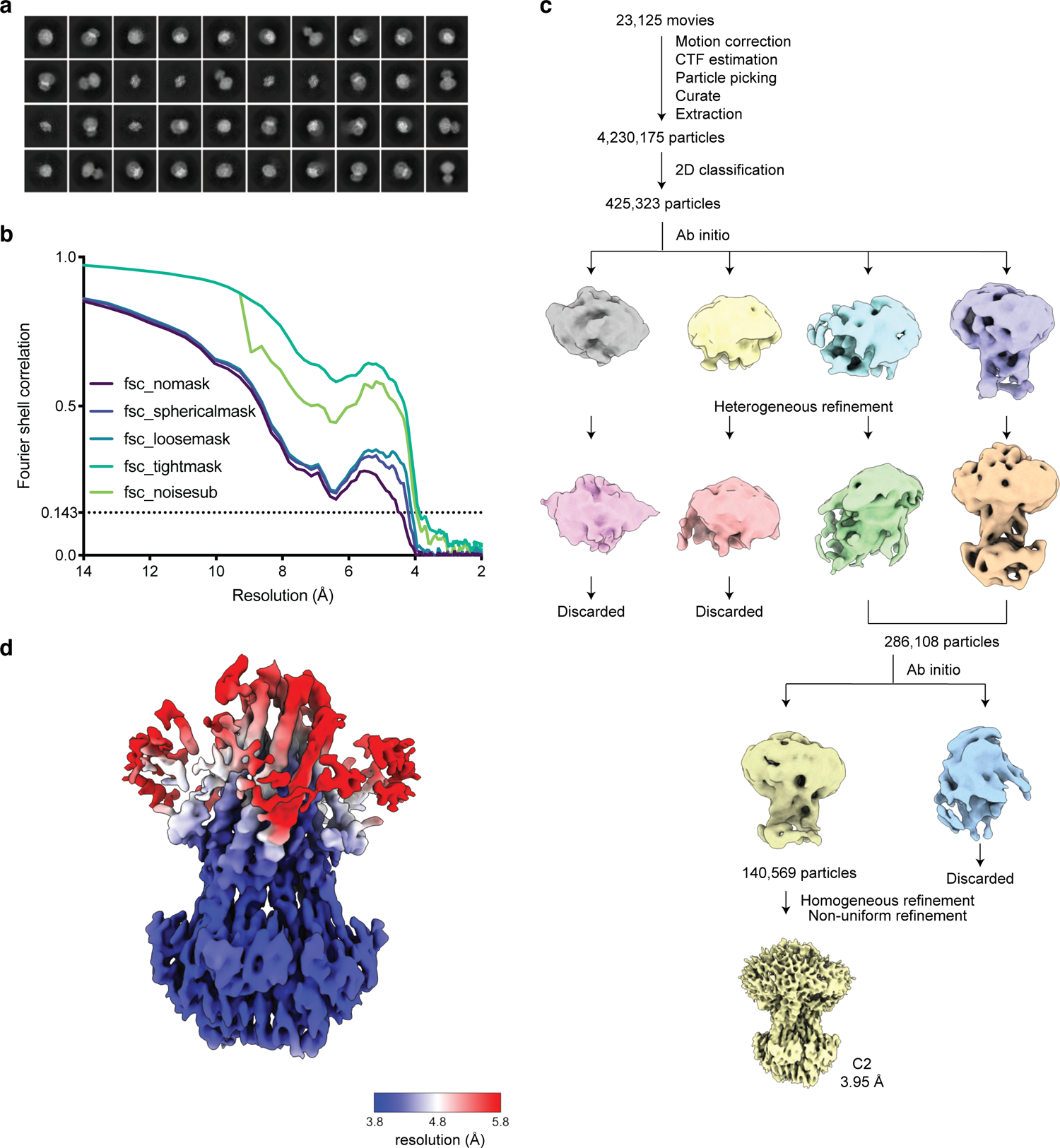
Single particle cryoEM structure of *At*Atm3 in the closed conformation. **a**) Examples of 2D classes, **b**) FSC curves showing the resolution estimate for the final reconstruction, **c**) workflow of single-particle data processing, and **d**) local resolution estimation of the *At*Atm3 closed conformation.

**Figure S7.**
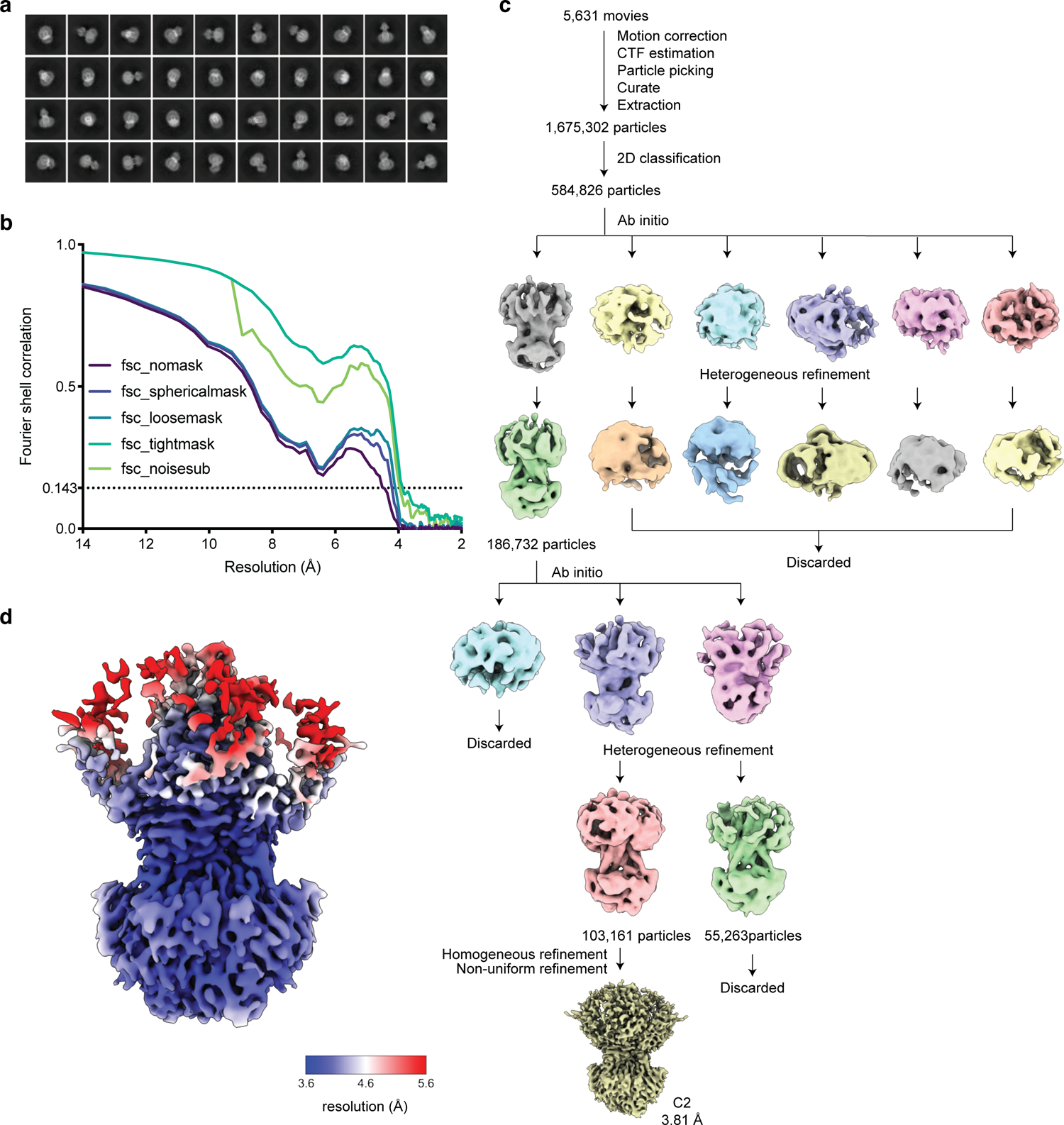
Single particle cryoEM structures of *At*Atm3 in the outward-facing conformation. **a**) Examples of 2D classes, **b**) FSC curves showing the resolution estimate for the final reconstruction, and **c**) workflow of single-particle data processing, and **d**) local resolution estimation of the *At*Atm3 outward-facing conformation.

**Figure S8.**
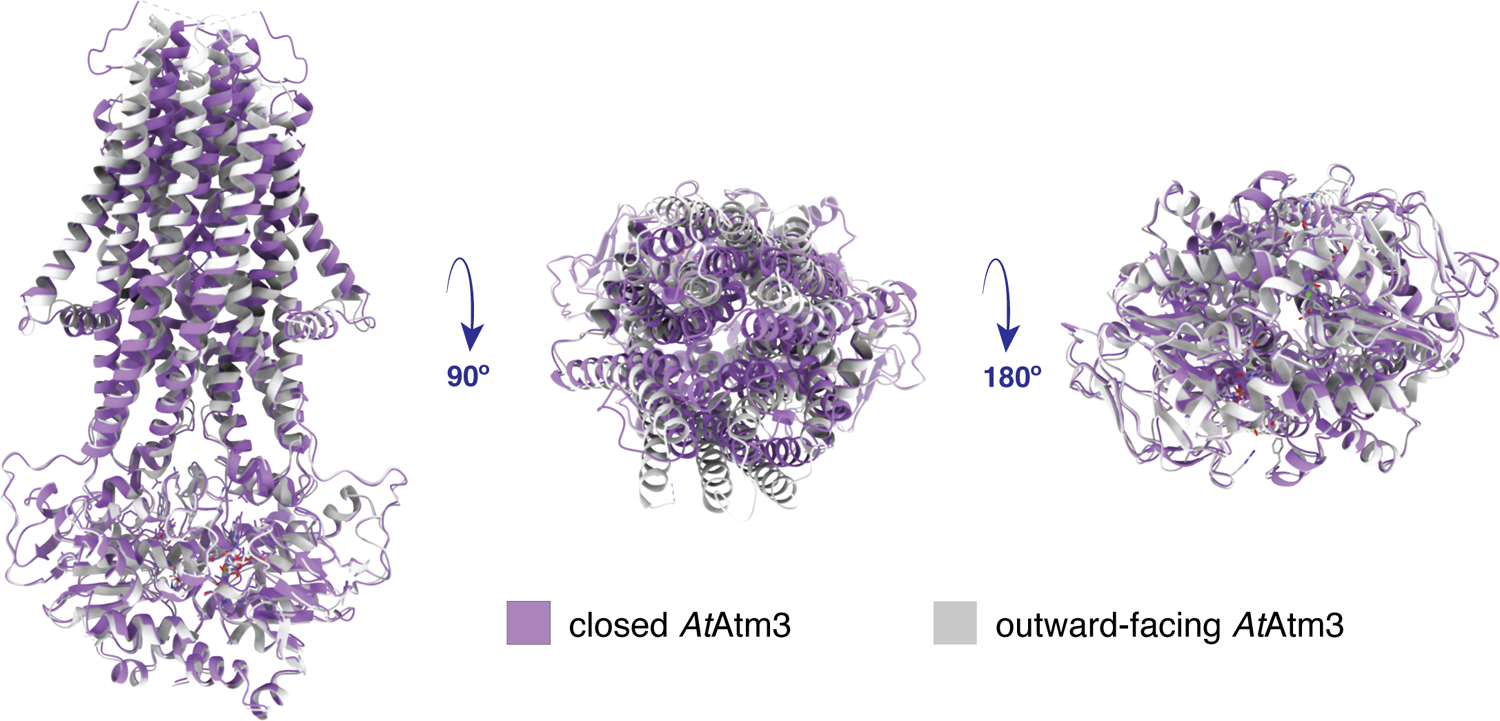
Structural alignment of *At*Atm3 in the closed and outward-facing conformation. Overall alignment of *At*Atm3 in the closed and outward-facing conformation with an overall RMSD of 1.7 Å.

**Figure S9.**
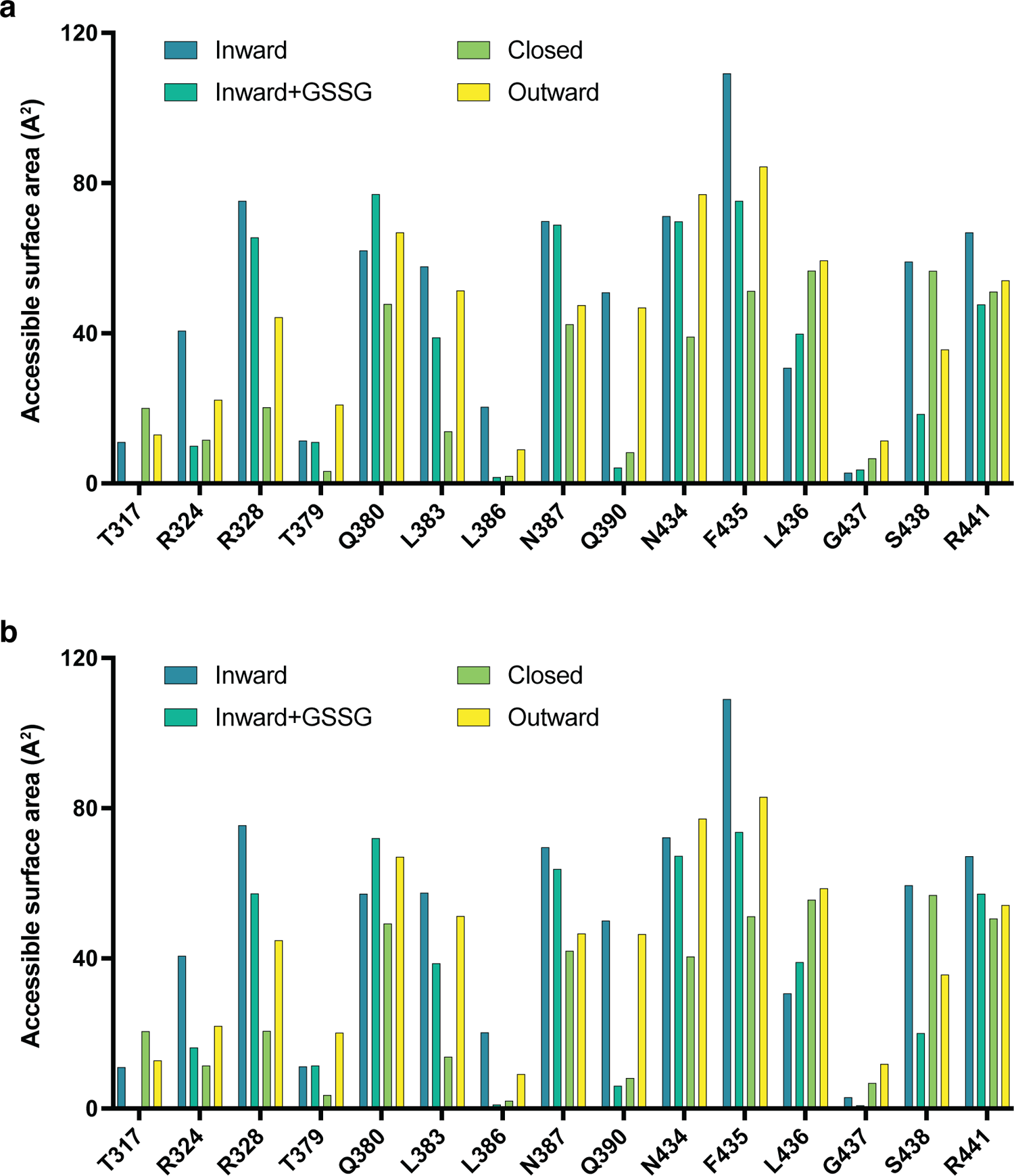
Accessible surface araa of binding site residues. Accessible surface area for GSSG binding pocket residues in **a**) chain A and **b**) chain B, calculated by Areaimol in CCP4 (Winn et al.2011).

**Figure S10.**
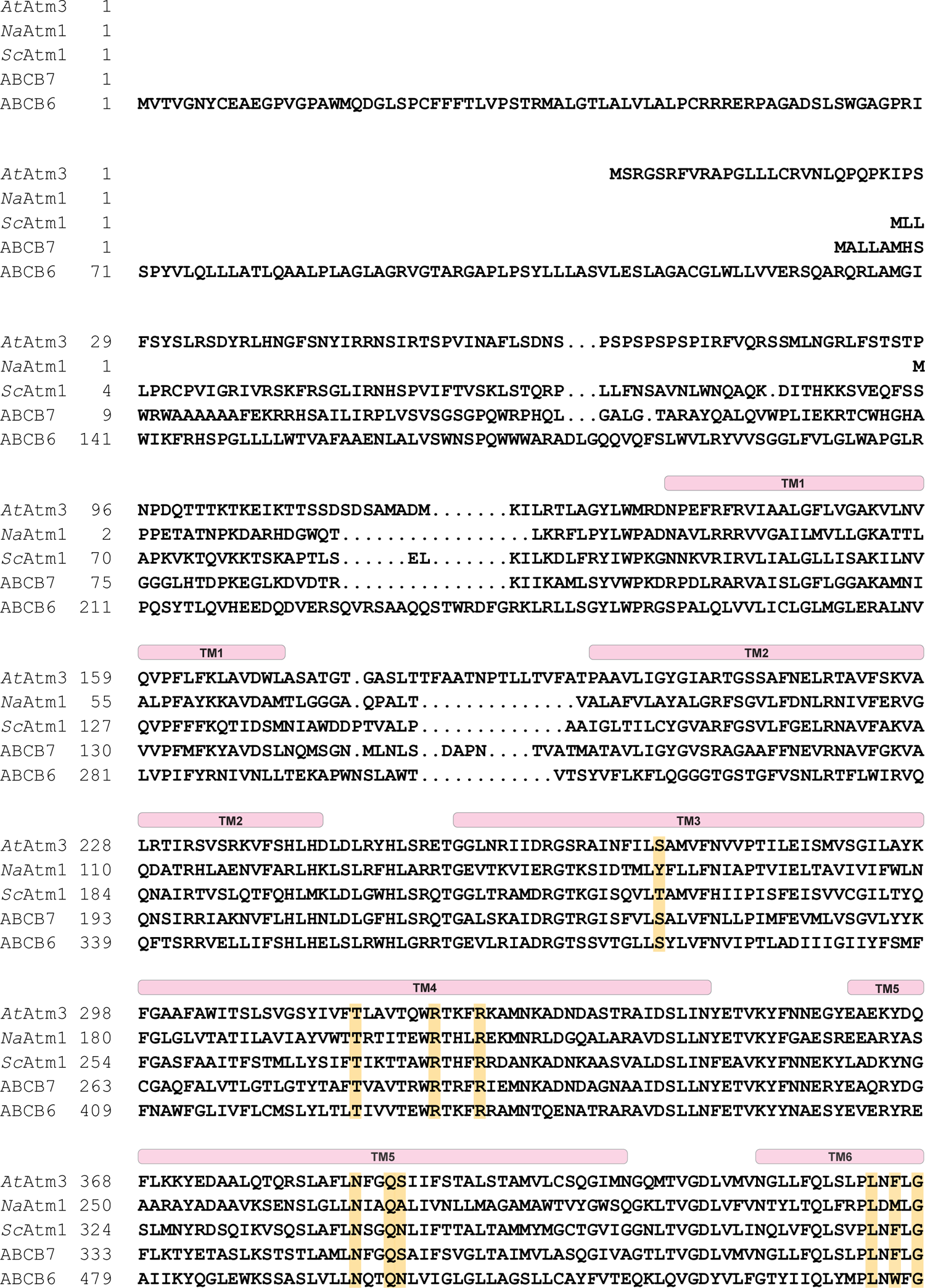
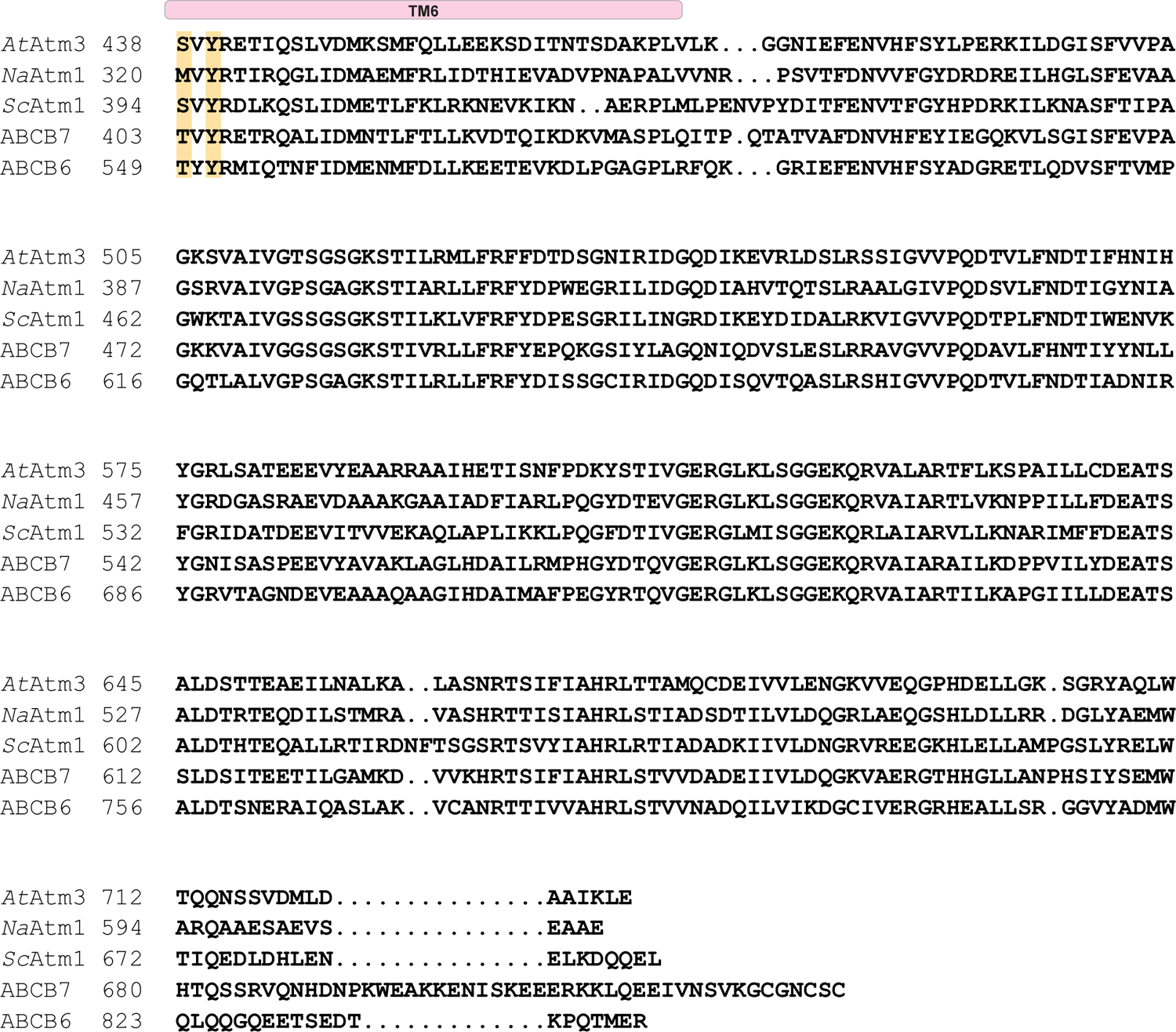
Sequence alignments of selected Atm family transporters. Sequence alignment generated by was generated using STRAP (http://www.bioinformatics.org/strap/). *At*: *Arabidopsis thaliana*; *Na*: *Novosphingobium aromaticivorans and Sc*: *Saccharomyces cerevisiae* (yeast). ABCB7 and ABCB6 are human ABC transporters. Positions of the six transmembrane helices are indicated above the sequence alignment. The key residues interacting with GSSG or GSH are highlighted in yellow. The substrate interacting residues are identified based on the structures of GSSG-bound *Na*Atm1 (PDB ID: 4mrs) and *At*Atm3 (PDB ID: 7n59), and the GSH-bound structure of *Sc*Atm1 (PDB ID: 4myh).

**Table S1.** Cryo-EM data collection, refinement and validation statistics

|  | Inward | Inward + GSSG | Closed | Outward |
| --- | --- | --- | --- | --- |
| **Data Collection and processing** |  |  |  |  |
| **Microscope** | Titan Krios at Caltech Cryo-EM facility | Titan Krios at Caltech Cryo-EM facility | Titan Krios at Stanford-SLAC CryoEM Center (S2C2) | Titan Krios at Caltech Cryo-EM facility |
| **Camera** | Gatan K3 | Gatan K3 | Falcon IV | Gatan K3 |
| **Magnification** | x105,000 | x105,000 | - | x105,000 |
| **Voltage (keV)** | 300 | 300 | 300 | 300 |
| **Exposure (e/Å<sup>2</sup>)** | 60 | 60 | 48 | 60 |
| **Pixel size (Å)** | 0.855 | 0.855 | 0.82 | 0.855 |
| **Defocus Range (um)** | - 1.0 to -3.0 | - 1.0 to -3.0 | - 1.5 to -2.1 | - 1.0 to -3.0 |
| **Initial Particle Image (no.)** | 4,608,600 | 2,510,131 | 4,230,175 | 1,675,302 |
| **Final Particle Image (no.)** | 157,762 | 259,020 | 140,569 | 103,161 |
| **Symmetry Imposed** | C2 | C2 | C2 | C2 |
| **Map Resolution (Å)** | 3.4 | 3.6 | 4.0 | 3.8 |
| **FSC Threshold** | 0.143 | 0.143 | 0.143 | 0.143 |
| **Map Resolution Range (Å)** | 3.2 - 4.1 | 3.5 - 4.0 | 3.9 - 4.2 | 3.9 - 4.3 |
| **Refinement** |  |  |  |  |
| **Initial Model Used** | PDB ID: 6pam | PDB ID: 6pam | PDB ID: 6par | PDB ID: 6par |
| **Model Resolution (Å)** | 3.4 | 3.57 | 3.95 | 3.81 |
| **FSC Threshold** | 0.143 | 0.143 | 0.143 | 0.143 |
| **Model composition** |  |  |  |  |
| **non-hydrogen atoms** | 9,326 | 9,238 | 9,254 | 9,122 |
| **protein residues** | 1,200 | 1,180 | 1,178 | 1,160 |
| **ligands** | - | GDS:1 | ADP: 2; MG: 2; VO<sub>4</sub>:2 | ADP: 2; MG: 2; VO<sub>4</sub>:2 |
| **Average B-factors (Å<sup>2</sup>)** |  |  |  |  |
| **protein** | 99.8 | 76.5 | 118.7 | 49.3 |
| **ligands** | - | 34.3 | 62.1 | 25.3 |
| **R.m.s. deviations** |  |  |  |  |
| **Bond length (Å)** | 0.004 | 0.002 | 0.003 | 0.002 |
| **Bond angles (°)** | 0.841 | 0.562 | 0.613 | 0.482 |
| **Validation** |  |  |  |  |
| **MolProbity score** | 1.82 | 1.41 | 1.60 | 1.41 |
| **Clashscore** | 14.6 | 6.9 | 11.8 | 6.3 |
| **Rotamer outliers** | 0 | 0.1 | 0.2 | 0.2 |
| **Ramachandran plot** |  |  |  |  |
| **Ramachandran favored (%)** | 97.2 | 97.9 | 98.0 | 97.7 |
| **Ramachandran allowed (%)** | 2.8 | 2.1 | 2.1 | 2.3 |
| **Ramachandran outliers (%)** | 0 | 0 | 0 | 0 |
| **PDB ID** | 7n58 | 7n59 | 7n5a | 7n5b |

**Table S2.** Principal component analysis. Calculated component 1 values are listed for different transporters. For heterologous transporters, transporter encoded in one polypeptide and transporters with different conformational states in one PDB file, “_A/B” is added at the end of each PDB to represent different half-transporters and/or different conformations.

| PDB ID | Component 1 (A<sup>2</sup>) | PDB ID | Component 1 (A<sup>2</sup>) |
| --- | --- | --- | --- |
| 2hyd | 0 | 5uj9_B | -93.27 |
| 3b5w | -252.01 | 5uja_A | -87.87 |
| 3b5x | -95.19 | 5uja_B | -63.67 |
| 3b5y | -0.09 | 5w81_A | -41.29 |
| 3b5z | 2.61 | 5w81_B | -45 |
| 3b60 | -0.59 | 6bhu_A | -25.63 |
| 3g5u_A | -112.24 | 6bhu_B | -23.02 |
| 3g5u_B | -52.24 | 6bpl | -84.93 |
| 3qf4_A | -115.57 | 6c0v_A | -10.35 |
| 3qf4_B | -109.08 | 6c0v_B | -19.67 |
| 3wme | -90.79 | 6fn4_A | -85.55 |
| 3zdq | -125.91 | 6fn4_B | -72.26 |
| 4ayt | -117.99 | 6hrc | 0.8 |
| 4ayw | -121.61 | 6msm | -36.8 |
| 4ayx | -133.59 | 6qee_A | -87.78 |
| 4f4c | -168.99 | 6qee_B | -84.19 |
| 4mrs | -47.43 | 6quz_A | -4.53 |
| 4myc | -111.84 | 6quz_B | 2.57 |
| 4pl0 | -21.98 | 6qv2_A | -3.93 |
| 4q4a_A | -101.87 | 6qv2_B | 1.11 |
| 4q4a_B | -110.01 | 6raf_A | -86.61 |
| 4ry2 | -68.98 | 6raf_B | -64.64 |
| 5c73 | -19.25 | 6rag_A | -107.09 |
| 5c76 | -187.37 | 6rag_B | -94.21 |
| 5c78 | -215.98 | 6rah_A | -9.19 |
| 5eg1 | -20.71 | 6rah_B | -12.44 |
| 5koy_A | -122.35 | 6rai_A | -15.64 |
| 5koy_B | -67.46 | 6rai_B | -27.8 |
| 5mkk_A | -97.26 | 6ral_A | -26.39 |
| 5mkk_B | -99 | 6ral_B | -35.32 |
| 5nbd | -122.85 | 6pam_A | -57.51 |
| 5ofp | -11.32 | 6pam_B | -73.71 |
| 5ofr | -20.86 | 6pan | -15.17 |
| 5ttp | -22.3 | 6pao | -17.01 |
| 5tv4 | -99.97 | 6paq | -15.42 |
| 5u1d_A | -129.2 | 6par | -29.28 |
| 5u1d_B | -93.49 | 6vqu | -131.4 |
| 5uak_A | -90.58 | 6vqt | -23.27 |
| 5uak_B | -83.07 | 7n58 | -128.9 |
| 5uar_A | -86.26 | 7n59 | -100.4 |
| 5uar_B | -69.55 | 7n5a | -25.82 |
| 5uj9_A | -137.22 | 7n5b | -15.71 |

